# Modelling the depth-dependent VASO and BOLD responses in human primary visual cortex

**DOI:** 10.1101/2021.05.07.443052

**Authors:** Atena Akbari, Saskia Bollmann, Tonima S Ali, Markus Barth

## Abstract

Functional magnetic resonance imaging (fMRI) using a blood-oxygenation-level-dependent (BOLD) contrast is a common method for studying human brain function non-invasively. Gradient-echo (GRE) BOLD is highly sensitive to the blood oxygenation change in blood vessels; however, the spatial signal specificity can be degraded due to signal leakage from activated lower layers to superficial layers in depth-dependent (also called laminar or layer-specific) fMRI. Alternatively, physiological variables such as cerebral blood volume using the VAscular-Space-Occupancy (VASO) contrast have shown higher spatial specificity compared to BOLD. To better understand the physiological mechanisms such as blood volume and oxygenation changes and to interpret the measured depth-dependent responses, models are needed which reflect vascular properties at this scale. For this purpose, we extended and modified the “cortical vascular model” previously developed to predict layer-specific BOLD signal changes in human primary visual cortex to also predict a layer-specific VASO response. To evaluate the model, we compared the predictions with experimental results of simultaneous VASO and BOLD measurements in a group of healthy participants. Fitting the model to our experimental data provided an estimate of CBV change in different vascular compartments upon neural activity. We found that stimulus-evoked CBV change mainly occurs in small arterioles, capillaries and intracortical arteries, and that the contribution from venules and ICVs is small. Our results confirm that VASO is less susceptible to large vessel effects compared to BOLD, as blood volume changes in intracortical arteries did not substantially affect the resulting depth-dependent VASO profiles, whereas depth-dependent BOLD profiles showed a bias towards signal contributions from intracortical veins.

## 1 Introduction

High-resolution functional magnetic resonance imaging (fMRI) offers the potential to measure depth-dependent hemodynamic responses, which can provide insights into cortical information processing and microcircuits of the human brain (Douglas and Martin, 2004; Lawrence et al., 2019; Stephan et al., 2019). Numerous studies have investigated the function of cortical layers using the blood-oxygenation-level-dependent (BOLD) contrast (Ogawa et al., 1990) in animals and humans (Aitken et al., 2020; Bollmann and Barth, 2020; Chen et al., 2013; de Hollander et al., 2021; Goense et al., 2012; Goense and Logothetis, 2006; Koopmans et al., 2010; Polimeni et al., 2010; Poplawsky et al., 2015; Ress et al., 2007; Self et al., 2017; Silva and Koretsky, 2002; van Dijk et al., 2020; Vizioli et al., 2020; Yu et al., 2014; Zaretskaya et al., 2020); for a brief history of the field see also Norris and Polimeni (2019). Despite the high sensitivity of this technique, it suffers from limited specificity due to signal leakage in draining veins carrying blood from (activated) lower layers to superficial layers and further to the pial veins (Duvernoy et al., 1981; Kim et al., 1994; Turner, 2002). This low specificity was the motivation to develop non-BOLD contrast mechanisms, such as cerebral-blood-volume (CBV) imaging, which is expected to be predominantly sensitive to hemodynamic responses in the microvasculature (Gagnon et al., 2015; Jin and Kim, 2006, 2008a; Kim and Kim, 2010, 2011a; Poplawsky et al., 2015; Silva et al., 2007; Vanzetta et al., 2005; Zhao et al., 2006).

A non-invasive method for CBV imaging is vascular-space-occupancy (VASO) (Lu et al., 2003), which takes advantage of the difference in blood and tissue *T*_1_ to image the tissue signal while the blood signal is nulled (Huber et al., 2014b; Lu et al., 2003). Since the development of this contrast and its translation to 7 Tesla (T), several studies in animals and humans have been conducted in the areas of method development (Beckett et al., 2019; Chai et al., 2019; Huber et al., 2015; Huber et al., 2016; Yu et al., 2014), analysis strategies (Huber et al., 2021; Polimeni et al., 2018), and applications to cognitive neuroscience (Finn et al., 2019; Huber et al., 2014a; Huber et al., 2017a; Kashyap et al., 2018; Oliveira et al., 2021b; Van Kerkoerle et al., 2017). However, to interpret the experimental results and account for both neural and vascular contributions to the fMRI signal, detailed models are required (Buxton et al., 2004). Several studies have modelled the BOLD response for both low- and high-resolution acquisitions (Baez-Yanez et al., 2020; Buxton et al., 2004; Buxton et al., 1998; Gagnon et al., 2015; Havlicek and Uludağ, 2020; Heinzle et al., 2016; Markuerkiaga et al., 2016; Uludağ et al., 2009). Recently, Genois et al. (2021) modelled BOLD and VASO signals using a vascular anatomical network (VAN) model (Boas et al., 2008) of the rat brain to investigate intra- and extra-vascular contributions to the BOLD signal and BOLD contribution to the VASO signal. However, no simulations of depth-dependent BOLD or VASO signals were provided in this study. Given the potential of VASO imaging for layer fMRI, we set out to model the depth-dependent VASO signal changes in human primary visual cortex (V1) employing a detailed model of the underlying macro- and micro-vasculature (Markuerkiaga et al., 2016). In this work, we extended and modified the “cortical vascular model” (Markuerkiaga et al., 2016) to simulate VASO responses in addition to BOLD responses at the laminar level. This model is based on histological observations in macaque primary visual cortex and considers various vascular features, such as vessel diameter, length, density, and distribution to simulate intra- and extra-vascular BOLD and VASO signals across cortical layers. We added intra-cortical (diving) arteries (ICAs) to the modelled region, as it is hypothesized that these play a role in the functional VASO response based on previous observations (Gagnon et al., 2015; Vanzetta et al., 2005). We also modified the artery to vein ratio defined in the model such that it is applicable to the human brain (Cassot et al., 2009; Schmid et al., 2019). To fit the predictions of the now extended model to experimental data, we performed simultaneous BOLD and VASO imaging in a group of healthy participants with sub-millimetre resolution at 7T. The model fitting then provided estimates of CBV and oxygenation changes in micro-vascular (arterioles, capillaries, venules) and macro-vascular (ICAs and intra-cortical veins (ICVs)) compartments at each cortical depth. Furthermore, we investigated the sensitivity of both VASO and BOLD contrasts to changes in the underlying physiological parameters, i.e., CBV and oxygenation.

To the best of our knowledge, this is the first study reporting depth-dependent experimental VASO profiles in the human primary visual cortex and the first depth-dependent VASO simulation study investigating the underlying physiological mechanism, such as the effect of baseline CBV on the resulting depth-dependent BOLD and VASO profiles. In the following sections, we briefly summarize the general structure of the previously developed cortical vascular model (2.1 and 2.2), and then describe the applied changes to simulate the VASO and BOLD responses (2.3 and 2.4).

## 2 Theory and Simulations

### 2.1 The cortical vascular model

The cortical vascular model developed by Markuerkiaga et al. (2016) simulates the steady-state BOLD response in a depth-dependent manner in human primary visual cortex. The model divides the brain vasculature into two groups: (i) the microvasculature forming a network of tangled, randomly oriented arterioles, capillaries, and venules called the *laminar network*, where the vessel distribution and blood volume varies as a function of depth, and (ii) The macrovasculature in the form of intra-cortical veins (ICVs) that drain the microvasculature towards the cortical surface. In the original version of the cortical vascular model (Markuerkiaga et al., 2016), the vessel distribution in the laminar network was 21% arterioles, 36% capillaries, and 43% venules. However, these values are based on the rat brain (Boas et al., 2008), and the artery-to-vein ratio should be reversed for the human brain (Cassot et al., 2009; Schmid et al., 2019). Thus, we assumed the following vessel density in the laminar network: 43% arterioles, 36% capillaries, and 21% venules. The model investigates the effect of the ascending veins on the BOLD signal by calculating the diameter, blood velocity, and mass flux of the ICVs in each layer. To simulate the VASO response, we added diving arteries to the modelled region, as several studies have shown that dilation mainly occurs in arteries and arterioles (Gagnon et al., 2015; Kim and Kim, 2011a; Vanzetta et al., 2005). To do so, a vascular unit centered on two adjacent principal veins (V3 and V4), surrounded by an arterial ring is modelled in this work with an artery-to-vein ratio of 2-to-1 (Cassot et al., 2009; Lauwers et al., 2008; Schmid et al., 2019). The diameter of this venous unit is around 0.75-1 mm. Considering the 2.5 mm thickness of V1 (Fischl and Dale, 2000), we simulated ten voxels with the size of 0.75×0.75×0.25 mm^3^, i.e. signal change of one venous unit at ten cortical depths. Figure 1A shows a schematic of the modelled intra-cortical arteries and veins following Duvernoy et al. (1981) in which vessels are categorized based on their diameter and penetration depth.

**Figure 1:**
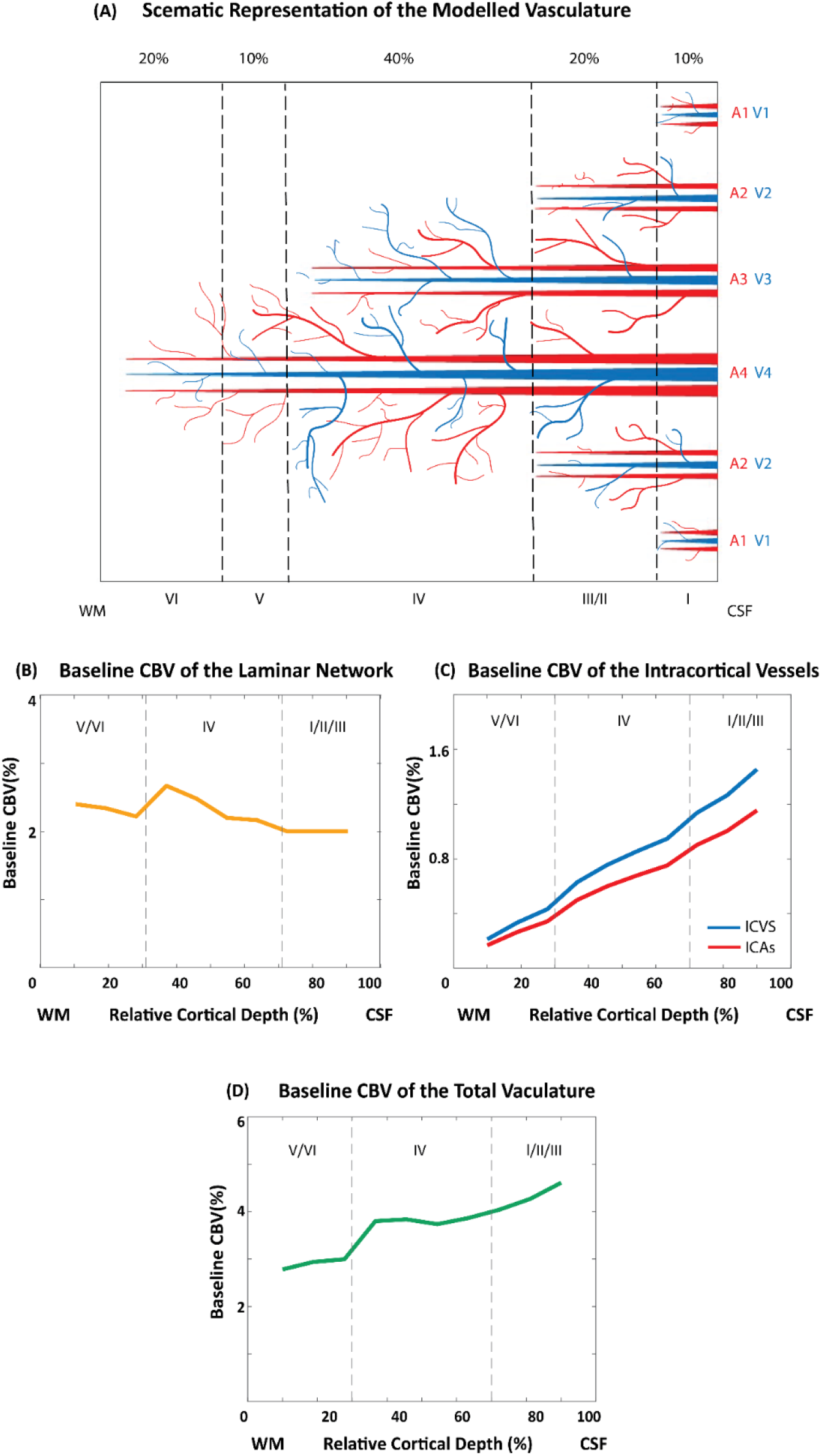
Vascular model of intra-cortical vessels and baseline cerebral blood volume of the different vascular compartments. A) Schematic of the vascular features of the primary visual cortex illustrating the 2 – 1 artery-to-vein ratio. B) Baseline blood volume of the laminar network (i.e., arterioles, capillaries, and venules) as a function of cortical depth following Weber et al. (2008). C) Estimated baseline blood volume of the intra-cortical vessels (ICAs and ICVs). D) Estimated baseline blood volume of the total vasculature.

The diameter of the intra-cortical vessels in each depth is calculated following the steps described in Markuerkiaga et al. (2016). In brief, based on the mass conservation law, the incoming mass flux (*p*) to the arteries should be equal to the outgoing flux from the veins in the modelled region at steady state. Similarly, the mass flux from each depth is the mass flux from within the laminar netwrok plus the mass flux in the macrovasculature from the previous layer. In general, the mass flux through vessels can be calculated as:

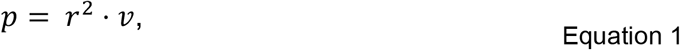

where *r* is the vessel radius and *v* is the blood velocity. We can rewrite this as:

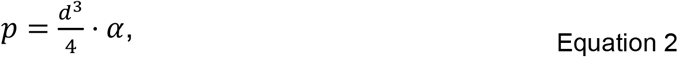

assuming a linear relationship between vessel diameter *d* and blood velocity (Zweifach and Lipowsky, 1977):

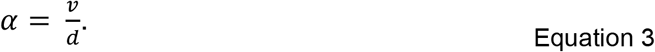

The mass flux *p* through a single capillary is calculated assuming *d* = 8 *μm* and *v* = 1.6 *mm/s* (Boas et al., 2008; Zweifach and Lipowsky, 1977). Then, *p* through ICVs and ICAs present in each layer is calculated starting from the layer closest to the white-matter (WM) border based on the number of capillaries in that layer:

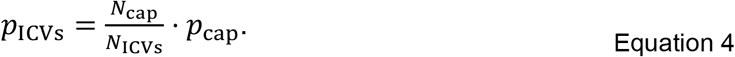

For the rest of the layers, *p* in each layer is the summation of the mass flux within that layer and the mass flux from the previous layer(s). The same calculation for ICVs applies for ICAs, but with twice the number of arteries (Schmid et al., 2019). Note that for this steady-state model, the direction of flow does not play a role when calculating the mass flux at each depth. The only relevant considerations are the mass conversation law, i.e., mass flux entering a layer should be equal to the mass flux that exits this layer, and the general layout of the model, i.e., blood from deeper layers flows through upper layers. Thus, the mass flux in each layer is equal to the additional mass flux (*p*_cap_) from the laminar network in that layer plus the mass flux in the preceding layers. Therefore, the calculation can be done in both bottom-up or top-down directions. The α values in pre- and post-capillaries compartments were calculated assuming *d* = 8 *μm* and *v* = 2 *mm/s* in post-capillary and *v* = 4 *mm/s* in the pre-capillary segment of the vasculature (Equation 3) (Zweifach and Lipowsky, 1977). Then, based on the mass flux for each vessel at each layer, the vessel diameter and the blood velocity of the macrovasculature can be calculated using Equation 2 and 3. Table 1 shows the estimated diameters of intra-cortical arteries and veins, which are in line with the values reported in Duvernoy et al. (1981).

**Table 1.** The average diameter (in µm) of intra-cortical arteries and veins in the modelled vascular unit centred on two intermediate-sized veins (V3 and V4) and surrounded by four intermediate-sized arteries (two A3 and two A4). For reference, the ICV diameters of group 1 to 4 reported in Duvernoy et al. (1981) range from 20 to 65 µm, and the diameter of the corresponding ICAs range from 10 to 40 µm.

| ***The Average Diameter ($\mu\text{m}$) of the Intracortical Vessels*** |  |  |  |  |  |  |  |  |
| --- | --- | --- | --- | --- | --- | --- | --- | --- |
| ***Vessel Type*** | ***V4*** | ***V3*** | ***V2*** | ***V1*** | ***A4*** | ***A3*** | ***A2*** | ***A1*** |
| ***Layer I*** | **69** | **53** | **32** | **20** | **43** | **33** | **20** | **13** |
| ***Layer II/III*** | **67** | **51** | **26** |  | **43** | **32** | **16** |  |
| ***Layer IV*** | **62** | **41** |  |  | **39** | **26** |  |  |
| ***Layer V*** | **56** |  |  |  | **35** |  |  |  |
| ***Layer VI*** | **44** |  |  |  | **28** |  |  |  |

The baseline blood volume of the laminar network taken from Weber et al. (2008) was interpolated to the number of voxels being simulated, resulting in 2-2.7% baseline CBV (Figure 1B). The intracortical baseline CBV is calculated as:

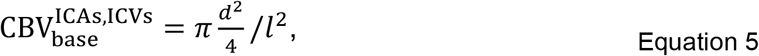

in which *l* is the simulated voxel length (0.75 mm) yielding a baseline CBV in ICVs ranging from 0.2-1.4% and a baseline CBV in ICAs ranging from 0.2-1.2% (Figure 1C). The average of the total baseline CBV of the modelled vasculature is 3.7 % (Figure 1D), where venous and arterial baseline CBV fractions are 57% and 43% of the total CBV, respectively. This is comparable to a PET study reporting arterial CBV fractions of 30 % in humans (An and Lin, 2002; Ito et al., 2005; Ito et al., 2001). This vascular model is then combined with the MR signal model (see section 2.2).

### 2.2 BOLD and VASO MR signal models

The MR signal model employed here is a steady state model contrasting signal levels at baseline and during activity (Markuerkiaga et al., 2016; Uludağ et al., 2009). At baseline, the total MR signal 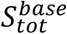 is the sum of the volume-weighted intra-(IV) and extra-vascular (EV) signal components (Buxton, 2009; Obata et al., 2004; Uludağ et al., 2009):

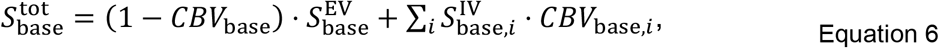

where CBV is the baseline blood volume, and *i* denotes different vascular compartments, i.e., arterioles, capillaries, venules, ICVs, and ICAs. In the following, we describe the intra- and extra-vascular BOLD and VASO signals when using a GRE readout at 7T.

The BOLD signal is approximated as a mono-exponential decay (Yablonskiy and Haacke, 1994), where *T*_*E*_ is the echo time, *S*_0_ the effective spin density at *T*_*E*_ = 0, and 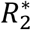 the transverse relaxation rate:

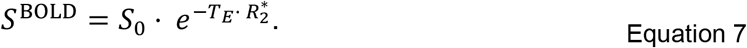

The transverse relaxation rate is the sum of the intrinsic 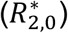 and hemoglobin (Hb)-induced transverse relaxation rates 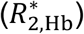:

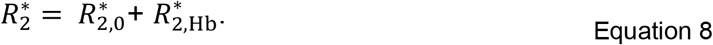

All intrinsic and Hb-induced 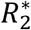 values used in this model (Blockley et al., 2008; Uludağ et al., 2009) are summarized in Table 2. Extra- and intra-vascular BOLD signals are estimated using their corresponding relaxation rates. In short, the Hb-induced extra-vascular relaxation rate is calculated according to the susceptibility-induced shift at the surface of the vessel depending on the oxygenation level (*Y*) (Uludağ et al., 2009). The intra-vascular 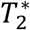 of the ICVs (intrinsic and Hb-induced) are very short at high field (7T and above). Therefore, the intra-vascular signal in veins approaches zero (Uludağ et al., 2009), and the main intra-vascular contribution comes from the arterial and capillary side of the vasculature.

**Table 2:**
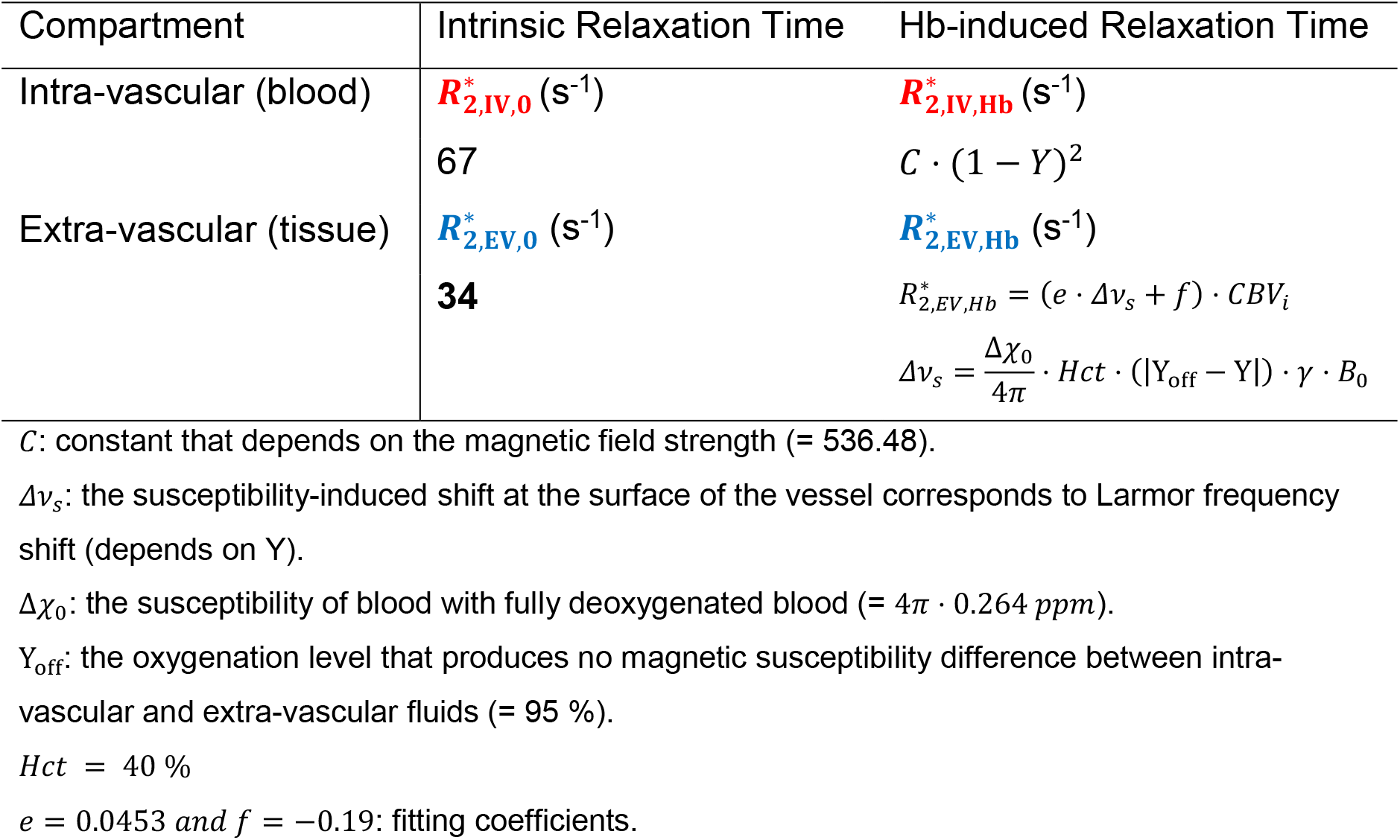
The intrinsic and Hb-induced intra- and extra-vascular transverse relaxation rates used in the BOLD signal model (Uludağ et al., 2009).

Following neural activity and changes in blood volume and oxygenation, the total MRI signal is:

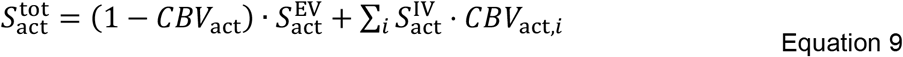

where *CBV*_*act*_ = *CBV*_*base*_ + Δ*CBV*_*abs*_ and Δ*CBV*_*abs*_ = Δ*CBV*_*rel*_ · *CBV*_*base*_. Note that Δ*CBV*_*abs*_ denotes the blood volume change upon activation in ml/100ml tissue commonly used (Huber et al., 2015; Lu et al., 2013) and Δ*CBV*_*rel*_ in % baseline blood volume, which is the definition implemented in the cortical vascular model. The increase in oxygenation is reflected in the shortening of the relaxation rates, which leads to increased extra- and intra-vascular signal levels. The BOLD signal change in percent (%) following neural activity can be described as the signal difference between baseline and activation, normalized to the baseline signal level:

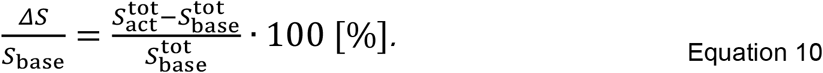

For VASO, assuming a perfect inversion pulse, signal change arises only from the extravascular component, as the intra-vascular signal is nulled with an inversion pulse. The steady state nulled tissue signal is (Huber et al., 2014b; Lu et al., 2003; Lu et al., 2013):

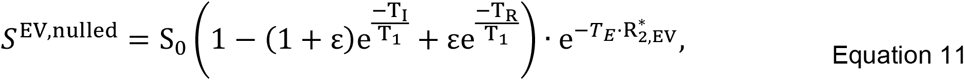

in which ε is the inversion efficiency (here assumed to be equal to 1), *T*_*I*_/*T*_1_/*T*_*R*_ are the blood nulling time, longitudinal relaxation time, and repetition time, respectively. At the time of the blood nulling, a BOLD signal contamination — the 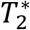-dependency — is still present and needs to be corrected. The dynamic division approach proposed by Huber et al. (2014b) removes the 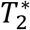-contribution from the VASO signal by dividing the “nulled” by the “non-nulled” signal, assuming equal extravascular BOLD contributions in both images and negligible intravascular BOLD signal:

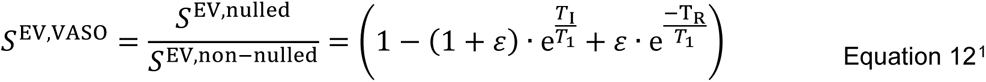

Then, the VASO signal during baseline, activity, and total VASO signal change can be derived from Equation 6, 9, and 10 by considering only the extra-vascular components:

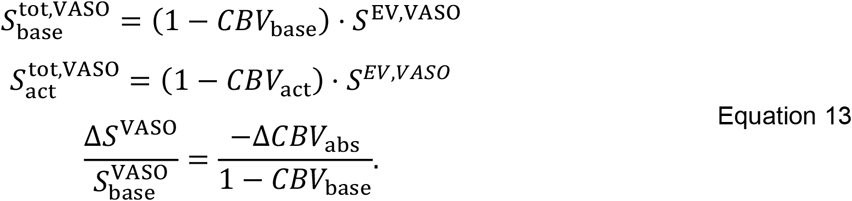

Thus, VASO signal changes are only a function of CBV change and baseline CBV, and independent of oxygenation changes.

### 2.3 Model Assumptions and Simulations

To simulate depth-dependent BOLD and VASO signal changes, the cortical vascular model outlined in section 2.1 requires ΔCBV and oxygenation values at baseline and activity for each depth and vascular compartment. Note that ΔCBV here denotes ΔCBV_rel_, which is given in percent of the baseline CBV, i.e., an increase of 100 % means that CBV during activation is twice as large as during baseline. Further, oxygenation is given in percent oxygen saturation, with 100 % oxygenation corresponding to fully oxygenated blood. The resulting BOLD and VASO profiles are presented in percent signal change following Equation 10 and 13.

To find the input values that best fit the empirical data (see Section 3.2) we simulated numerous profiles for a wide range of input parameters (Table 3), and then calculated the root-mean-squared-error (RMSE) for each simulated profile with the experimental result. The minimum RMSE value indicates the highest similarity between simulated and measured depth-dependent responses. To investigate the effect of input parameters on the resulting depth-dependent profiles, we also plotted the range of profiles obtained with RMSEs that are 20 % higher than the minimum RMSE and extracted the corresponding input values for ΔCBV and oxygenation. For reference, we have included the original values used in Markuerkiaga et al. (2016) in square brackets in Table 3.

**Table 3:** Range of the model parameters for simulating VASO and BOLD depth-dependent responses. The values in brackets refer to the values used in the original vascular model (Markuerkiaga et al., 2016). *Y*_*base*_ and *Y*_*act*_ are the blood oxygenation at baseline and activation, and Δ*CBV*_*mid*_ refers to the CBV change in middle layer. In the laminar network, the CBV change in middle layers is 1.5 times higher than in deep and superficial layers. In ICAs and ICVs, ΔCBV in middle and upper layers is 1.5 times higher than the change in deep layers.

| Vascular Compartments CBV and Oxygenation |  |  |  |  |  |
| --- | --- | --- | --- | --- | --- |
|  | arterioles | capillaries | venules | ICVs | ICAs |
| $Y_{base}$ | 95 %<br>[95 %] | 85 %<br>[85 %] | 60- 75* %<br>[60 %] | 60-75* %<br>[60 %] | 95 %<br>-- |
| $Y_{act}$ | 100 %<br>[100 %] | 95 %<br>[95 %] | 75-90<sup>†</sup> %<br>[70 %] | 75-90<sup>†</sup> %<br>[70 %] | 100 %<br>-- |
| $\Delta CBV_{mid}$ | 0-90 %<br>[16.6 %] | 0-90 %<br>[16.6 %] | 0-90 %<br>[16.6 %] | **0-90 %**<br>0 % | 0-90 %<br>-- |

\*Corresponds to $R_{2,IV}^* = 100 - 153 \text{ sec}^{-1}$
<sup>†</sup>Corresponds to $R_{2,IV}^* = 72 - 100 \text{ sec}^{-1}$

To account for partial volume effects across layers and provide a better comparison between simulated and measured depth-dependent responses (Markuerkiaga et al., 2016), we applied a smoothing kernel (Koopmans et al., 2011) to the simulated profiles. The resulting zero-padded edges of the laminar profiles were thus excluded from the RMSE estimation and all simulated profiles, and only the central eight data points contributed.

In the original version of the cortical vascular model, Markuerkiaga et al. (2016) assumed the same increase in CBV of 16.6 % (Griffeth et al., 2015) in each compartment of the laminar network across all depths. Using these input values, the resulting depth-dependent VASO signal change (see Figure 8) did not exhibit the characteristic peak in the middle layers observed in our and other VASO experiments (Huber et al., 2014a). We therefore assumed a non-uniform activation strength across the layers (Equation 14). For the VASO simulations, we assumed in the laminar network a 1.5 times higher CBV increase in middle cortical layers (*IC*) compared with deep (*C* and *CI*) and superficial (*I* and *II/III*) layers (Zhao et al., 2006). In ICAs and ICVs, we assumed higher CBV change in middle and upper cortical layers compared with deep layers (see Figure 2):

**Figure 2:**
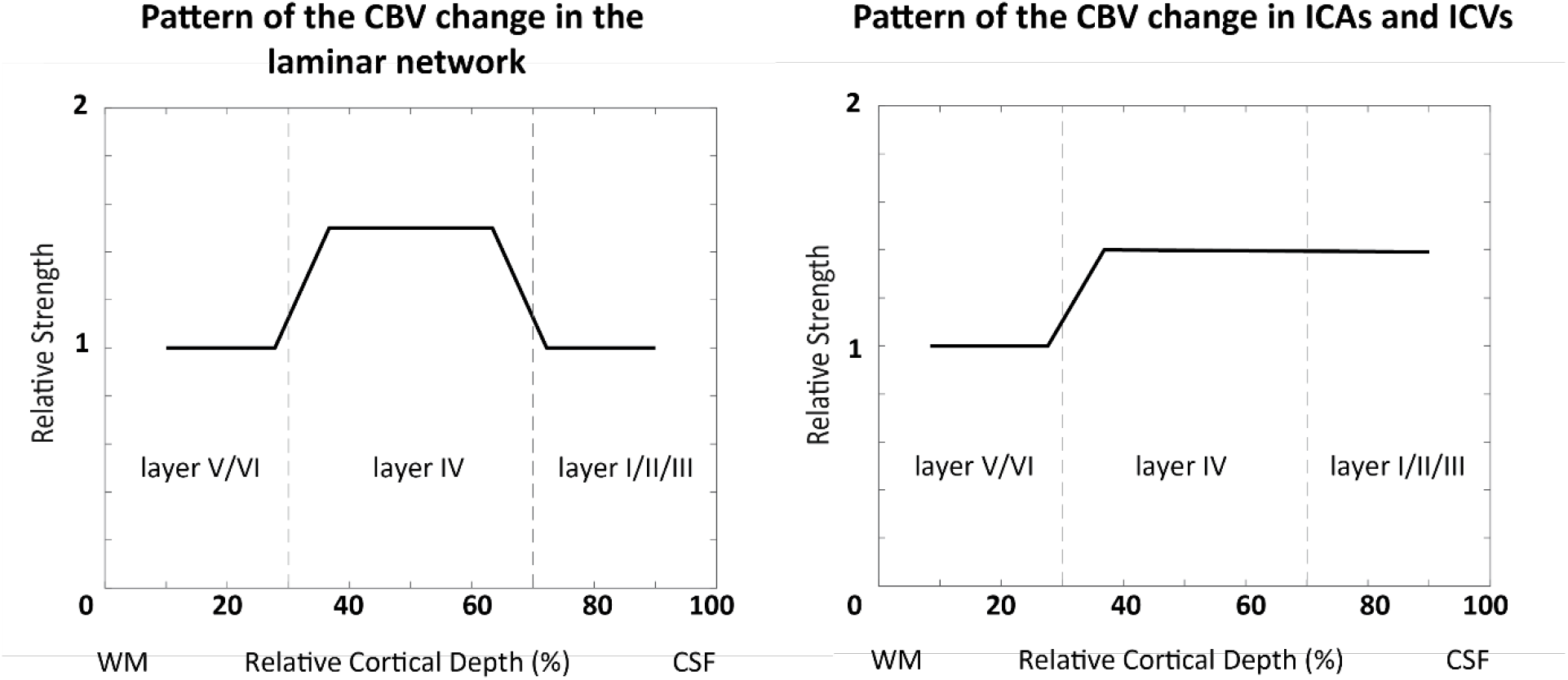
The pattern of the CBV change in the laminar network (left), and ICAs and ICVs (right) across the layers used in our simulations. According to the result shown in Figure 3 of Zhao et al. (2006), we assumed a higher CBV change in middle cortical layers by a factor of 1.5 in the laminar network. In ICAs and ICVs, the assumption is that the CBV change in deep layers is 2/3 of the change in middle and superficial layers.

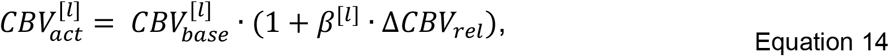

in which *β* refers to the activation strength and *β* = 1.5 for middle depths, *β* = 1 for superficial and deep cortical depths (Figure 2), and *l* denotes the cortical layers. We then simulated VASO profiles using this ratio for a range of ΔCBV values of 0 - 90 % for ICAs, arterioles, capillaries, venules, and ICVs (Table 3). In the original version of the cortical vascular model (Markuerkiaga et al., 2016), ICVs were considered not to dilate as shown by Hillman et al. (2007). However, other studies in cats (Kim and Kim, 2011b), mice (Takano et al., 2006), and humans (Chen and Pike, 2009; Stefanovic and Pike, 2005) observed dilation in ICVs, in particular for long stimulus duration. Therefore, we allowed this parameter to vary as well.

For the BOLD simulations, we used the ΔCBV values of the best fit from the VASO experiment, and instead varied oxygenation values between 60-75 % at baseline and 75-90 % at activation in the venules and ICVs (see Table 3). Following Markuerkiaga et al. (2016) and Uludağ et al. (2009), we assumed fixed oxygen saturation in ICAs, arterioles and capillaries at baseline and activation as outlined in Table 3, but varied the oxygen saturation of the venules and ICVs at baseline and activation to find the best fit.

### 2.4 The effect of depth-dependent vs constant baseline CBV on the simulated profiles

To investigate the effect of variation in baseline CBV across depths, we performed the same simulation and fitting procedures, but assumed a constant baseline CBV across depths in the laminar network. We used a baseline CBV value of 2.3 % across layers, which is the average of the depth-dependent baseline CBV (Weber et al., 2008, Figure 1B). Figure 3 shows the baseline CBV of the laminar network, the baseline CBV in ICVs and ICAs (model output), and the total CBV of the vasculature (average ∼3.7 %) for this scenario. The aim of these simulations was to investigate which impact the pattern of the baseline CBV (constant vs depth-dependent) has on the simulated BOLD and VASO responses.

**Figure 3:**
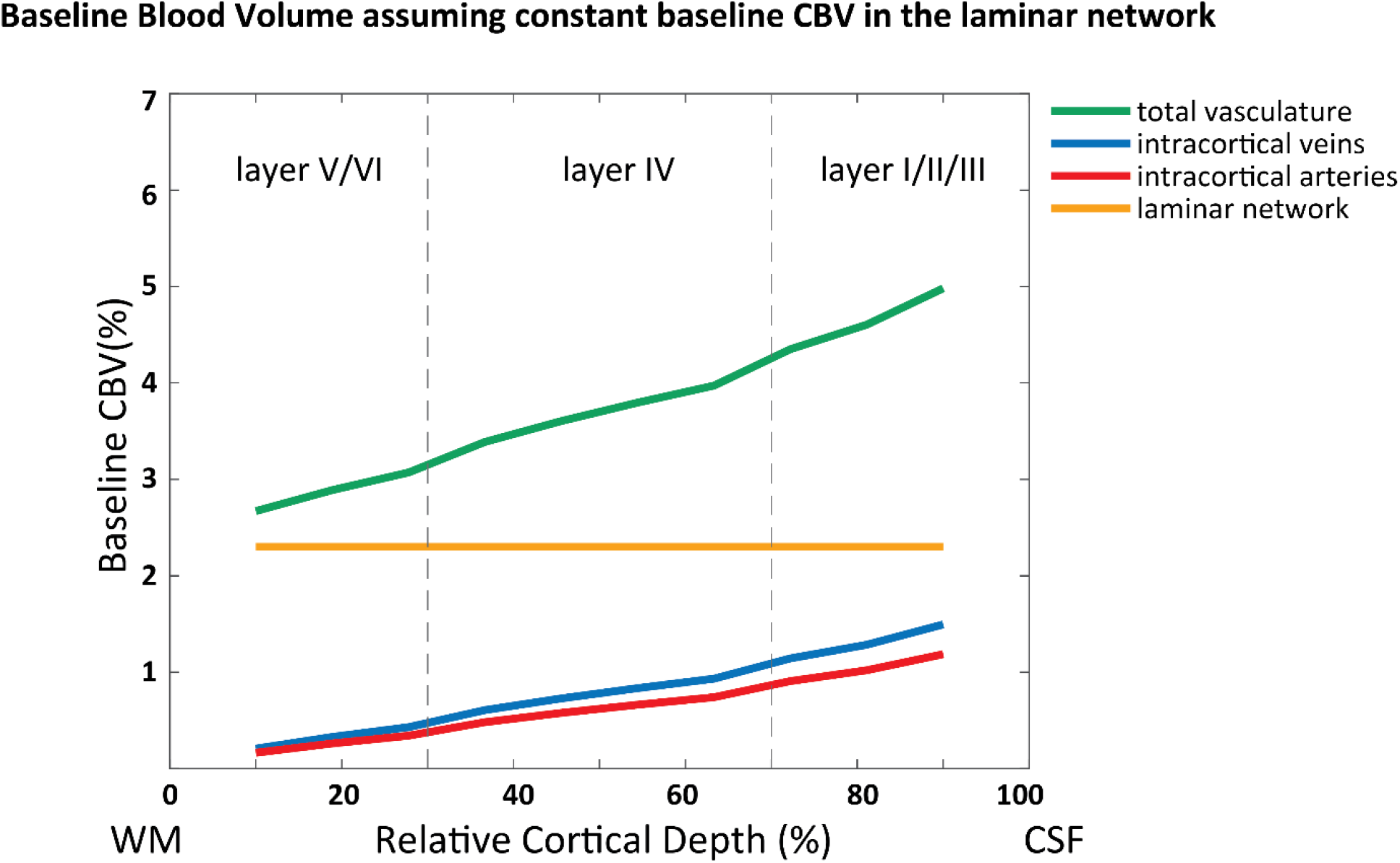
Baseline blood volume of the laminar network (i.e., arterioles, capillaries, venules), intra-cortical arteries, veins, and total vasculature in the “constant” baseline CBV scenario. We used the average of the baseline CBV in laminar network (2.3 %) reported in Weber et al. (2008).

## 3 Experimental Methods

### 3.1 Model Implementation

The cortical vascular model was implemented in MATLAB (2018b, The MathWorks, Inc.). The code is available on gitlab (https://gitlab.com/AtenaAkbari/cortical-vascular-model) including the original version used in Markuerkiaga et al. (2016) (branch: originalCode), the implementation used for an earlier version of this work presented at the ISMRM 2020 in which intra-cortical arteries were not yet added (Akbari et al., 2020) (branch: vasoSignal), and the implementation used in this manuscript (branch: master).

### 3.2 Image Acquisition

Imaging was performed on a 7T whole-body MR scanner (Siemens Healthcare, Erlangen, Germany), with a maximum gradient strength of 70 mT/m and a slew rate of 200 mT/m/s. A single-channel Tx and 32-channel Rx head coil array (Nova Medical, Wilmington, MA, USA) was used for radiofrequency transmission and signal reception. The slice-selective slab-inversion (SS-SI) VASO sequence (Huber et al., 2014b) was employed to scan ten healthy participants (2 females and 8 males; age range 19-32 years) after giving written informed consent according to the approval of the institutional ethics committee. For each subject, BOLD and VASO images were acquired in an interleaved fashion in three runs with 400 volumes in each run (15 minutes total acquisition time per run). The sequence parameters were: *T*_*R*_ = 4.5*s, T*_*E*_ = 25 *ms, T*_*I*_ = 1100 *ms*, GRAPPA (Griswold et al., 2002) acceleration factor = 3, isotropic voxel size = 0.8 *mm*^3^, number of slices = 26, partial Fourier in the phase encoding direction = 6*/*8, in combination with a 3D EPI readout (Poser et al., 2010). The blood-nulling time was chosen based on the assumed value of blood *T*_1_ = 2100 *ms* following earlier VASO studies at 7T (Huber et al., 2015; Huber et al., 2016; Zhang et al., 2013). The visual stimulus consisted of 15 ON- and OFF-blocks with 30 s duration each. During the ON condition, a flashing black and white noise pattern was presented, and a fixation cross was the OFF condition of the stimulus (Polimeni et al., 2005). The imaging slices were positioned and oriented such that the center of the imaging slab was aligned with the center of the calcarine sulcus, the part of the striate cortex with the highest vascular density in layer *IC* of V1 (Duvernoy et al., 1981). Whole-brain MP2RAGE images (Marques et al., 2010; O’Brien et al., 2014) were acquired with an isotropic resolution of 0.75 mm for each participant in the same session as the functional imaging.

### 3.3 Image Analysis

The first volume of each contrast was discarded to ensure *T*_1_ effects were at equilibrium. Dynamic division was then performed to account for the BOLD-contamination (Huber et al., 2014b). The BOLD and BOLD-corrected VASO images were motion corrected using SPM12 (Wellcome Department, UK). Activation maps were estimated with the GLM analysis in SPM with no spatial smoothing. Data from three participants were discarded due to excessive motion, i.e., volume-to-volume displacement of more than one voxel size. Voxels with t-values above 2.3 corresponding to an uncorrected significance level of *p* < 0.01 were identified as the activated regions for both BOLD and BOLD-corrected VASO images.

For the layer analysis, we followed the steps outlined in Huber et al. (2014b): The *T*_1_ − *EPI* images of each subject were used for WM/GM and GM/CSF boundary delineation, and a region of interest (ROI) was manually defined such that the activated regions in the calcarine sulcus from both contrasts were included (see Figure 4). Then, this ROI was used to create ten equi-volume layers (Waehnert et al., 2014) using the open-source LAYNII package (Huber et al., 2021) and extract depth-dependent BOLD and VASO responses. The average of the percentage signal change extracted from each layer forms the layer profile in both VASO and BOLD contrast. The mean and standard error of the mean were calculated across participants, and average BOLD and VASO responses across the seven participants were used as a reference when evaluating the model predictions. Note that cortical layers in these analyses refer to a group of voxels obtained by dividing the ROI into 10 equi-volume layers and do not refer to the histological cortical layers. In the next section, we will first present the imaging results, and then introduce the simulations that fit these best. Further, we will present the simulated profiles with an RMSE 20 % higher than the minimum RMSE, and the ΔCBV and oxygenation values corresponding to these profiles.

**Figure 4:**
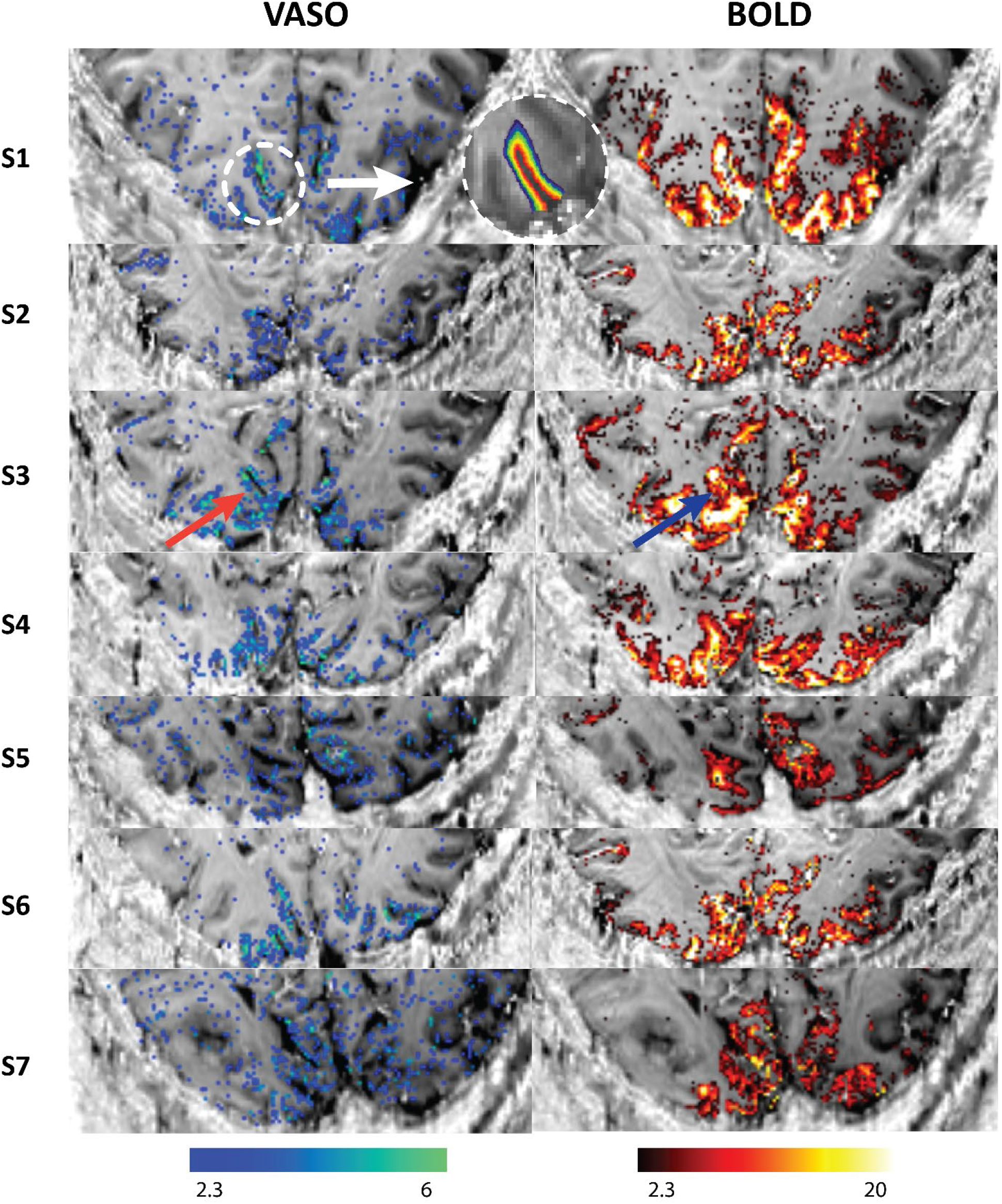
VASO and BOLD statistical activation maps of all participants in our study using the SS-SI VASO sequence (Huber et al., 2014b) with an isotropic resolution of 0.8 mm^3^. The activation maps are overlaid on *T*_1_ − *EPI* images of each subject. The VASO contrast is more confined to GM while BOLD shows higher activity near surface (indicated with the red and blue arrows). An example of the region-of-interest (ROI) in V1 for the layer analysis is shown above. Ten equi-volume layers were extracted from GM in the *T*_1_ − *EPI* images to calculate the mean signal change in each layer.

## 4 Results

### 4.1 Imaging

The BOLD and VASO activation maps of all seven participants included in this study are shown in Figure 4. We observed overall higher t-values for the BOLD contrast compared to the VASO contrast. Further, highest t-values for BOLD are located at the cortical surface and within various sulci. In contrast, most of the VASO response is confined to the grey matter. An example of the ROI placed on V1 to extract the 10 equi-volume layers and estimate the depth-dependent profiles is shown for one subject in Figure 4.

The depth-dependent BOLD and VASO signal changes for each participant as well as the mean and standard error of the mean of these profiles are shown in Figure 5. On average, we observed a mean signal change of 6 % for BOLD and 1 % for VASO, evidence for the larger effect size of the BOLD contrast. In agreement with previous studies, BOLD signal change peaks at the cortical surface (Fracasso et al., 2018; Koopmans et al., 2010; Olman et al., 2012; Polimeni et al., 2010) while the VASO signal change has its maximum in the middle cortical layers (Huber et al., 2014a).

**Figure 5:**
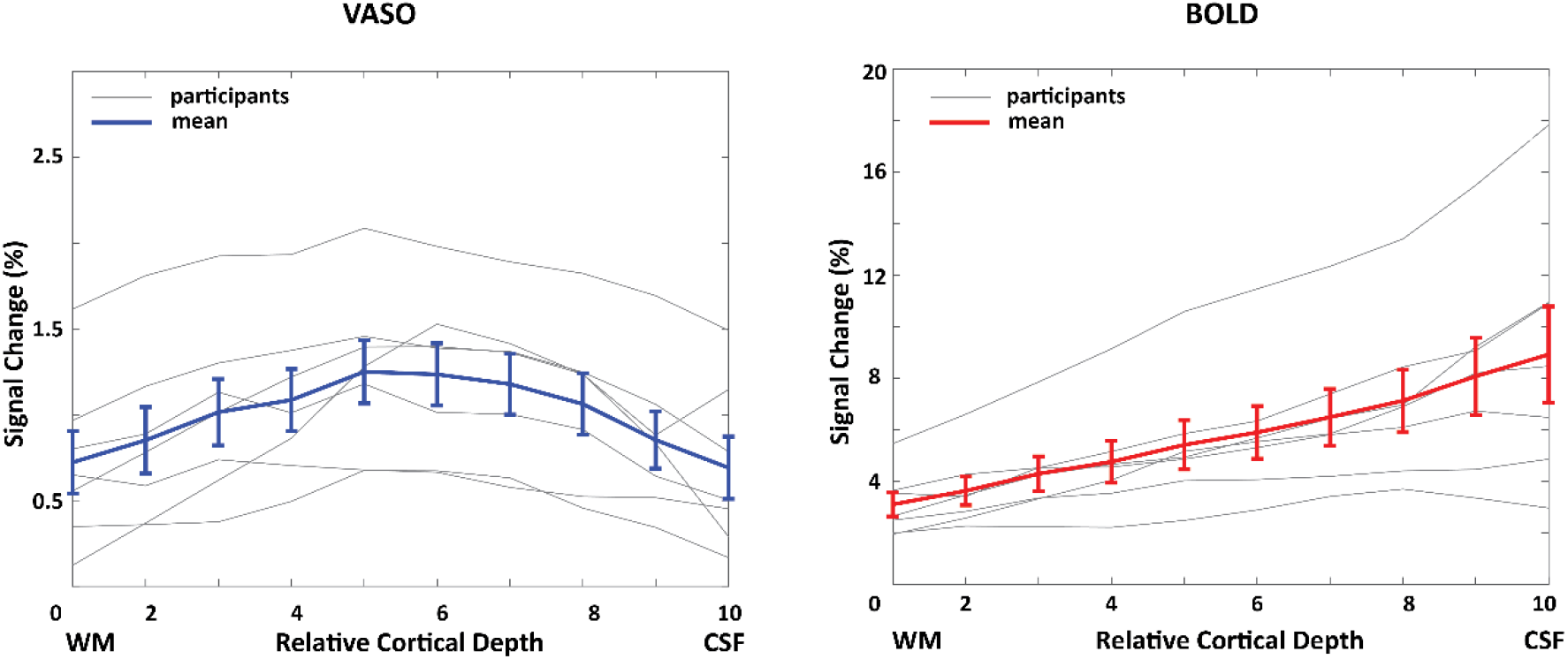
Depth-dependent VASO and BOLD signal changes (%) in human V1 for each individual participant (gray) and averaged across all participants (blue and red). Note that the profiles plotted here are the average of the percentage signal change extracted from each layer. The error bars in this and all following graphs refer to the standard error of the mean across all participants.

### 4.2 Simulations

#### 4.2.1 Depth-dependent BOLD and VASO profiles

The simulated VASO profile with the best fit using a depth-dependent baseline CBV in the laminar network (Figure 1) is shown in Figure 6, and corresponding CBV changes are shown in Table 4. We estimated highest CBV changes in arterioles and capillaries (56 %) and in ICAs (21 %), and small CBV changes in venules (2 %) and ICVs (5 %). Using a Grubb value (Grubb et al., 1974) of 0.35, the corresponding CBF change upon activation in middle layers would be 58 % in ICAs, 82 % in arterioles and capillaries, 25 % in venules, and 35 % in ICVs. For BOLD, the simulation with the best-fit yields *Y*_*base*_ = 70 % and *Y*_*act*_ = 90 % in venules and ICVs. To investigate the sensitivity of the model to the choice of input parameters, the shaded area in Figure 6 illustrates the range of profiles with an RMSE up to 20 % higher than the minimum RMSE. The resulting profiles remain predominantly within the standard error of the measured profiles.

**Table 4:** CBV changes in vascular compartments corresponding to the best-fit shown in Figure 6, i.e. assuming a depth-dependent baseline CBV. The minimum and maximum of the estimated CBV change in the 20 % RMSE regime (i.e., the minimum RMSE + 20 % of the minimum RMSE shown as the shaded area in Figure 6) are shown in brackets.

| **CBV Change in Vascular Compartments** |  |  |  |  |
| --- | --- | --- | --- | --- |
| **Cortical depth** | **Arterioles & capillaries** | **venules** | **ICAs** | **ICVs** |
| **Deep (V, VI)** | **37 %**<br>[21 % 41 %] | **1 %**<br>[0 % 49 %] | **14 %**<br>[0 % 42 %] | **3 %**<br>[0 % 42 %] |
| **Middle (IV)** | **56 %**<br>[32 % 62 %] | **2 %**<br>[0 % 73 %] | **21 %**<br>[0 % 63 %] | **5 %**<br>[0 % 63 %] |
| **Superficial (I, II/III)** | **37 %**<br>[32 % 62 %] | **1 %**<br>[0 % 49 %] | **21 %**<br>[0 % 63 %] | **5 %**<br>[0 % 63 %] |

**Figure 6:**
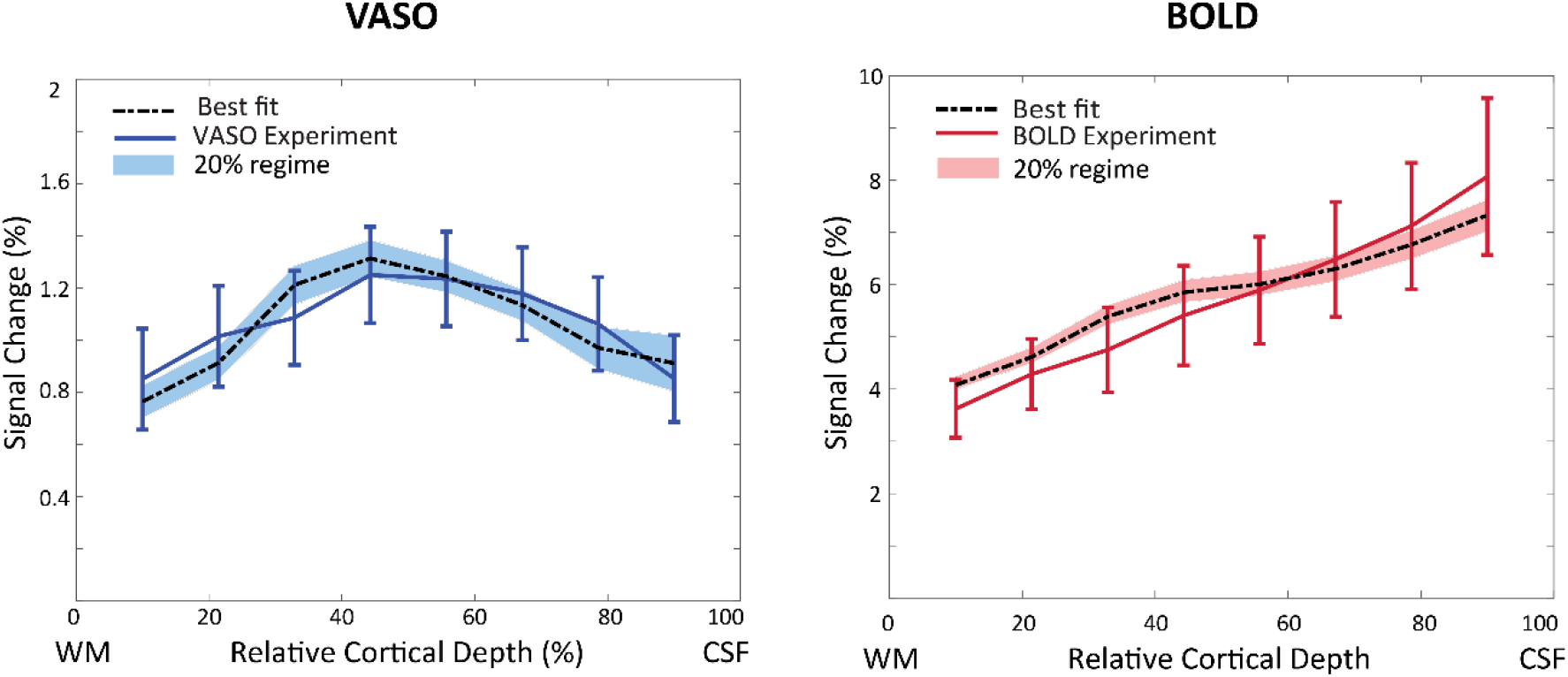
The measured VASO (left) and BOLD (right) profiles and the simulated profiles with the lowest RMSE (black line) assuming a depth-dependent baseline CBV (Figure 1). Shaded area shows the VASO and BOLD simulated profiles with RMSE 20 % higher than the minimum RMSE.

#### 4.2.2 The effect of constant baseline CBV on the simulated profiles

Figure 7 illustrates the simulation results assuming a constant baseline CBV across depths (Figure 3). This assumption produces a better fit for both VASO and BOLD profiles and a smaller 20 %-RMSE regime (shaded area). Note, however, that in the laminar network we are still assuming a 1.5 times higher CBV change in middle layers than in superficial and deep layers (Figure 2, Equation 14). Again, highest CBV change was estimated in arterioles and capillaries (65 % in middle layers), and small CBV changes in all other compartments (Table 5). For BOLD, the best fit estimates 71 % and 90 % oxygen saturation at baseline and activation, respectively.

**Table 5:** CBV changes in vascular compartments corresponding to the best fit shown in Figure 7, i.e., assuming a constant baseline CBV of 2.3 % in the laminar network. The minimum and maximum of the estimated CBV change in the 20 % RMSE regime (i.e., the minimum RMSE+20 % of the minimum RMSE shown as the shaded area in Figure 7) are shown in brackets.

| **CBV Change in Vascular Compartments** |  |  |  |  |
| --- | --- | --- | --- | --- |
| **Cortical depth** | **Arterioles & capillaries** | **venules** | **ICAs** | **ICVs** |
| **Deep (V, VI)** | **43 %** | **0 %** | **2 %** | **1 %** |
|  | **[28 % 45 %]** | **[0 % 49 %]** | **[0 % 10 %]** | **[0 % 10 %]** |
| **Middle (IV)** | **65 %** | **0 %** | **3 %** | **2 %** |
|  | **[42 % 68 %]** | **[0 % 73 %]** | **[0 % 15 %]** | **[0 % 15 %]** |
| **Superficial (I, II/III)** | **43 %** | **0 %** | **3 %** | **2 %** |
|  | **[28 % 45 %]** | **[0 % 49 %]** | **[0 % 15 %]** | **[0 % 15 %]** |

**Figure 7:**
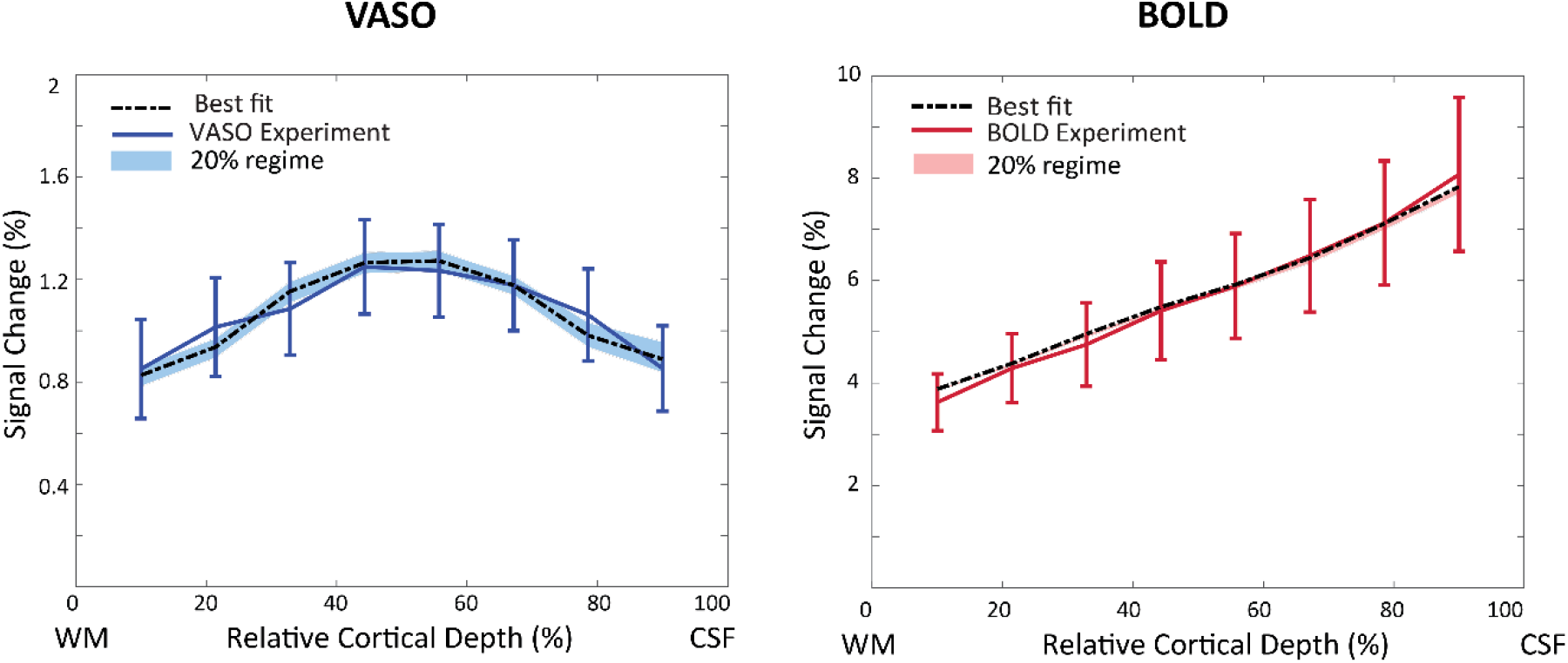
The measured and simulated VASO and BOLD profiles assuming a constant baseline CBV of 2.3 % in the laminar network. The shaded areas show the RMSEs that are 20 % higher than the minimum RMSE. The high agreement between BOLD simulation and measurement resulted in a low minimum RMSE, and a correspondingly very narrow 20 % RMSE regime.

#### 4.2.3 The effect of constant relative CBV changes across depths

The simulation results of equal activation strength across layers (i.e., *β* = 1 for all layers in Equation 14 such that all layers are activated equally) are shown in Figure 8. Note that this equal activation strength across depths was the assumption in the original implementation (Markuerkiaga et al., 2016). However, under this assumption, the VASO profile is much flatter and does not show the expected higher CBV change in middle layers (Goense et al., 2012; Poplawsky and Kim, 2014; Zhao et al., 2006). In contrast, this scenario had little impact on the predicted BOLD profile that showed comparable characteristics and similar fitting performance, but the experimental VASO profile could not be reproduced. The best fit estimates 48 % CBV change in arterioles and capillaries, 3 % in venules, 12 % in ICAs, and 9 % in ICVs in middle layers. The corresponding oxygenation levels at baseline and activation are 65 % and 86 %, respectively.

**Figure 8:**
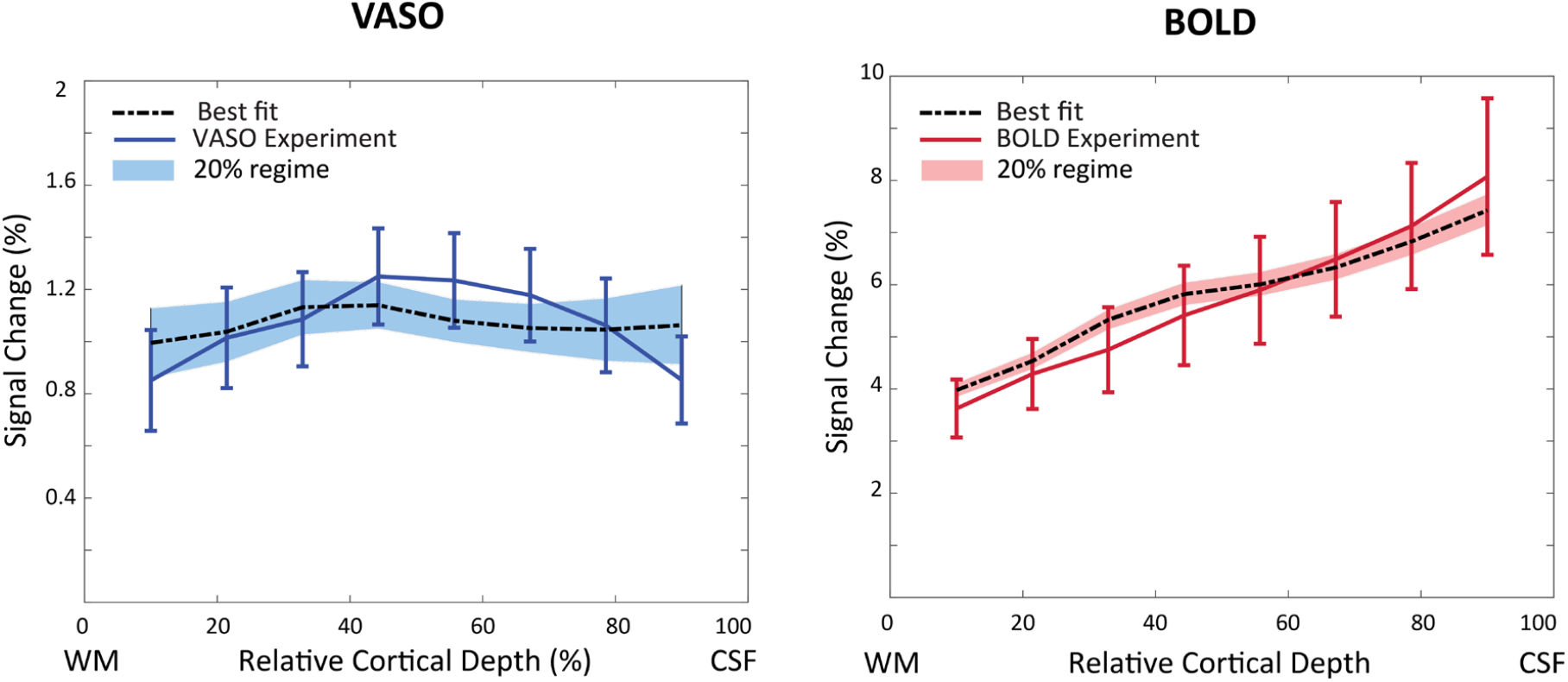
The measured and simulated VASO and BOLD profiles assuming a depth-dependent baseline CBV, but constant activation strength across depths. The simulated VASO profile deviates considerably from the measured response, whereas the fit to the BOLD data is comparable to the scenario with depth-dependent non-uniform activation strength.

## 5 Discussion and Conclusion

In this study, we extended and modified the cortical vascular model (Markuerkiaga et al., 2016) to simulate depth-dependent VASO signal changes in addition to BOLD signal changes and added intra-cortical arteries to the modelled area for a full description of CBV changes in the intra-cortical vasculature. With our simulations, we found that stimulus evoked CBV changes are dominant in small arterioles and capillaries at 56 % and in ICAs at 21 %, and that the contribution of venules and ICVs is small at 2 % and 5 %, respectively. These estimates of higher arterial CBV change are in line with previous studies, e.g. in mice with 60 s stimulus duration (Takano et al., 2006), cats with 40 s stimulus duration (Kim and Kim, 2010, 2011b) and humans with 30 s or longer stimulus duration (Chen and Pike, 2009, 2010; Huber et al., 2014a). Chen and Pike (2009, 2010) also showed small CBV changes in veins with 180 s and 96 s stimuli, respectively. In a simulation study using the extended Windkessel model (Mandeville et al., 1999), Barrett et al. (2012) estimated dominant CBV change in arteries (64.8 %) and smaller CBV change in veins (21.5 %) with 30 s stimulus duration. In addition, it is also expected that the VASO contrast is overall less sensitive to CBV changes in venules and veins due to water exchange in the capillaries (Huber et al., 2014b; Jin and Kim, 2008b; Lu et al., 2003). In summary, the cortical vascular model allows to estimate and compare BOLD and VASO signal changes in various conditions and resolves the contributions of different vascular compartments to the fMRI signal.

The inclusion of ICAs allowed us to investigate the sensitivity of the VASO signal to upstream CBV changes. Although we found a relatively large CBV change of 21 % in ICAs, the measured and simulated profiles did not show the pial bias that is commonly found in BOLD profiles (Barth and Norris, 2007; Kim et al., 1994; Turner, 2002). This might be due to the different contrast mechanisms of VASO and BOLD, where the VASO signal is linearly proportional to CBV (Equation 13). Together with the low baseline blood volume in ICAs (Figure 1C), even a relatively large CBV change in ICAs might thus only have a limited impact on the resulting VASO profile. In contrast, the extravascular signal contributions around venules and ICVs presumably amplify the effect of oxygenation changes in these vessels on the BOLD signal (Equation 7 and Table 2). Thus, the measured and simulated BOLD profiles are heavily skewed towards the signal stemming from ICVs. In conclusion, our results confirm that the VASO contrast is less susceptible to large vessel effects compared to BOLD (Huber et al., 2014b; Huber et al., 2017b; Lu et al., 2003; Lu et al., 2013). As the model fits the measured BOLD response well, we didn’t investigate the effect of the pial veins on the simulated laminar response. However, Markuerkiaga et al. (2021) added the pial vessels to the modelled area, and removed the contribution of downstream vasculature in laminar GRE-BOLD response using a deconvolution approach. This resulted in more spatially specific laminar BOLD responses that no longer show the bias towards the cortical surface.

In general, we found that a wide range of CBV changes in the different micro- and macro-vascular compartments can result in similar depth-dependent profiles (Table 4); indicating potential challenges when aiming to invert the measured profiles. These variations are illustrated in more detail using RMSE plots (see Figures S3 and S4 in the Supplementary Materials), where we observed high differentiability between macro- and micro-vascular compartments, but considerable interchangeability within each vascular class, such that for example a wide range of ΔCBV combinations in ICVs and ICAs can results in comparable VASO profiles, and thus comparable RMSE values. Similarly, a change in oxygenation of 20 % provides the best fit for BOLD, but a wide range of oxygenation levels at baseline and a corresponding value during activity are possible.

To investigate the influence of the assumed baseline CBV on the simulated profiles, we compared the effect of assuming either a depth-dependent or a constant baseline CBV in the laminar network and found that a constant baseline CBV across depths provides a better fit to both BOLD and VASO profiles. However, this observation might be driven by uncertainties in defining an accurate depth-dependent baseline CBV, namely (i) the baseline CBV used here was derived from *ex vivo* macaque data (Weber et al., 2008), (ii) the WM/GM and GM/CSF boundary definition was manually performed in EPI image space introducing potential inaccuracies, and (iii) the depth-dependent profiles were averaged across participants. Consequently, an *average*, i.e. constant baseline might perform better in this case.

A slight divergence—but within the standard error of the mean—remains between simulated and measured data. Notably, the so-called “bump” (Chen et al., 2013; Havlicek and Uludağ, 2020; Huber et al., 2017a; Koopmans et al., 2010) is visible in the simulated BOLD profile but not the imaging data. Preliminary investigations show that this feature seems to be mainly driven by the baseline CBV in this model (Figure 6 vs. Figure 7, see also Figure S1 in the Supplementary Materials), but in general is expected to have mixed neuronal and vascular origin (Havlicek and Uludağ, 2020). Note that this feature was also not evident in numerous studies using comparable acquisition and analysis techniques (de Hollander et al., 2021; De Martino et al., 2013; Fracasso et al., 2018; Klein et al., 2018; Marquardt et al., 2020; Polimeni et al., 2010).

The various parameters used to build the cortical vascular model such as blood velocity, vessel diameter, and baseline blood volume in capillaries were taken from previous research in rats, cats, rabbits, and macaques (Markuerkiaga et al., 2016; Weber et al., 2008; Zweifach and Lipowsky, 1977), which might impose uncertainties on the estimated CBV changes and simulated profiles. However, we took into account the inter-species differences in vascular structure and density by reversing the arterioles-to-venules ratio defined in the VAN model (Boas et al., 2008), as the arterial density in the human brain is higher than its venous density in contrast to the ratio in the rat brain (Cassot et al., 2009; Schmid et al., 2019). Here, we assumed an artery-to-vein ratio of 2-to-1 (Cassot et al., 2009; Schmid et al., 2019), but larger artery-to-vein ratios of 2.58 (Adams et al., 2015) and smaller artery-to-vein ratios of 1.6 (Weber et al., 2008) have also been reported in the literature for macaque brain. Further, we noticed that for certain parameter combinations the derived vessel diameters and blood velocities in the ICAs and ICVs can easily contradict previous reports that intracortical arteries have smaller diameter (Duvernoy et al., 1981) and faster blood velocities (Zweifach and Lipowsky, 1977). Thus, while the cortical vascular model aims for a detailed description of the underlying micro-and macro-vasculature and its influence on the MR signal, many uncertainties in the specific parameter choices remain. One example includes the dilation profile of the ICAs across cortical depths, where we assumed a higher CBV change in middle and superficial layers (Figure 2). However, another possible scenario could be an equal activation strength in ICAs across cortical depths (see Figure S5 and Figure S6 of the Supplementary Materials), which results in similar depth-dependent VASO and BOLD profiles, but different estimated CBV changes. Additionally, the inter-individual variability in these parameters remains unknown, but may potentially have a large effect on the individual profiles given the many studies showing significant differences in hemodynamic responses between participants (Aguirre et al., 1998; Duann et al., 2002; Handwerker et al., 2004; Light et al., 1993). Consequently, a more detailed understanding of the relative impact of each of these parameters needs to be developed, in combination with auxiliary image acquisitions that measure relevant underlying parameters.

The experimental results show similar profiles as expected from previous research (Huber et al., 2013; Huber et al., 2016; Jin and Kim, 2006, 2008b; Koopmans et al., 2010). To ensure highest contrast-to-noise ratio when comparing with the simulations, we have averaged the responses across participants. We extracted percent signal change values using a GLM, assuming the same hemodynamic response for all cortical layers. Although each layer has a unique HRF (Petridou and Siero, 2017), we expect a negligible bias in the estimated signal change due to the very long stimulus time employed here. There is also evidence of the dependency of blood *T*_1_ on hematocrit levels (Dobre et al., 2007) affecting the blood nulling time, though the effect can be considered negligible.

The cortical vascular model used here presents a simplification of vascular anatomical networks (VAN) (Boas et al., 2008; Gagnon et al., 2015; Genois et al., 2020), but employs more details in the mirco- and macro-vasculature than the fully invertible model developed by Havlicek and Uludağ (2020). Thus, it is uniquely suited to translate new insights from detailed VAN models developed in rat to the dynamic laminar models used to fit human data. As exemplified in this work using changes in CBV, the impact of each parameter on the resulting laminar profiles can be assessed individually, to then inform the choice of acquisition, potential vascular biases, and the need for auxiliary information. Next, the vascular anatomical model can be extended to other cortical areas characterized by different vascular properties such as primary motor cortex (Huber et al., 2017a; Oliveira et al., 2021a), primary somatosensory cortex (Shih et al., 2013; Silva and Koretsky, 2002), dorsolateral-prefrontal cortex (Finn et al., 2019), which are currently under active investigation using laminar fMRI to help to understand the vascular and neural signal contributions. In addition, the potential of higher spatial resolution can be explored with the model. For example, inspired by the study of Huber et al. (2015) who measured CBV responses in the macaque brain at 500 µm resolution and observed a double-peak pattern, i.e., local maxima on both sides of the stria of Gennari, we simulated a depth-dependent VASO profile but without applying any smoothing kernel. Interestingly, the model prediction indeed features a pattern that resembles a double peak corresponding to higher local CBV changes on both sides of stria of Gennari (Huber et al., 2015) (see Figure S7 in the Supplementary Materials).

In summary, we acquired BOLD and VASO laminar responses in human V1 at 7T and simulated these responses using the cortical vascular model. To the best of our knowledge, this is the first study to acquire the laminar BOLD and VASO profiles in addition to simulating these responses in the human primary visual cortex. By fitting the model to our experimental results, we obtained an estimate of CBV change in all vascular compartments upon neural activity. Our simulation results show that stimulus evoked CBV change is dominant in small arterioles and capillaries followed by ICAs, and the contribution of venules and ICVs in total CBV change is small when the stimulus is relatively long (∼30 sec). Our results also suggest that the large vessel bias is less prominent in VASO contrast compared with BOLD, as the BOLD signal relationship with the oxygenation change is exponential, but VASO depends on the CBV change linearly.

## 6 Declaration of interests

None.

## 7 Acknowledgement

We acknowledge the helpful discussions with and support from Laurentius (Renzo) Huber, Jonathan Polimeni, and Irati Markuerkiaga. We would also like the thank the anonymous reviewers for valuable feedback on the manuscript. We thank Aiman Al-Najjar and Nicole Atcheson for help with data collection. This work was supported by the NHMRC (grant APP1117020) and the NIH (grant R01-MH111419). MB acknowledges funding from ARC Future Fellowship grant FT140100865. AA acknowledges support through the University of Queensland Research Training Program Scholarship. We also acknowledge the facilities and scientific and technical assistance of the National Imaging Facility (NIF), a National Collaborative Research Infrastructure Strategy (NCRIS) at the Centre for Advanced Imaging, the University of Queensland.

## Footnotes

1 Note that 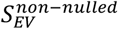 corresponds to the extravascular BOLD signal. During the BOLD correction of the experimental data, we assume the intravascular BOLD signal to be negligible following Huber et al. (2014), and Genois et al. (2021).

## Notes

### Competing Interest Statement

The authors have declared no competing interest.

## References

Adams, D.L., Piserchia, V., Economides, J.R., Horton, J.C., 2015. Vascular supply of the cerebral cortex is specialized for cell layers but not columns. Cerebral Cortex 25, 3673–3681. https://doi.org/10.1093/cercor/bhu221.

Aguirre, G.K., Zarahn, E., D’Esposito, M., 1998. The variability of human, BOLD hemodynamic responses. Neuroimage. 8, 360–369. https://doi.org/10.1006/nimg.1998.0369.

Aitken, F., Menelaou, G., Warrington, O., Koolschijn, R.S., Corbin, N., Callaghan, M.F., Kok, P., 2020. Prior expectations evoke stimulus-specific activity in the deep layers of the primary visual cortex. PLos biology. 18, e3001023. https://doi.org/10.1371/journal.pbio.3001023.

Akbari, A., Bollmann, S., Ali, T., Barth, M., 2020. Modelling the Laminar VASO Signal Change in Human V1 at 7T. 28th Annual Meeting of the International Society for Magnetic Resonance Imaging in Medicine.

An, H., Lin, W., 2002. Cerebral venous and arterial blood volumes can be estimated separately in humans using magnetic resonance imaging. Magnetic Mesonance in Medicine. 48, 583–588. https://doi.org/10.1002/mrm.10257.

Baez-Yanez, M.G., Siero, J.C., Petridou, N., 2020. A statistical 3D model of the human cortical vasculature to compute the hemodynamic fingerprint of the BOLD fMRI signal. bioRxiv. https://doi.org/10.1101/2020.10.05.326512.

Barrett, M.J., Tawhai, M.H., Suresh, V., 2012. Arteries dominate volume changes during brief functional hyperemia: evidence from mathematical modelling. Neuroimage. 62, 482–492. https://doi.org/10.1016/j.neuroimage.2012.05.005.

Barth, M., Norris, D., 2007. Very high-resolution three-dimensional functional MRI of the human visual cortex with elimination of large venous vessels. NMR in Biomedicine. 20, 477–484. https://doi.org/10.1002/nbm.1158.

Beckett, A.J., Dadakova, T., Townsend, J., Huber, L., Park, S., Feinberg, D.A., 2019. Comparison of BOLD and CBV using 3D EPI and 3D GRASE for cortical layer fMRI at 7T. BioRxiv., 778142.

Blockley, N., Jiang, L., Gardener, A., Ludman, C., Francis, S., Gowland, P., 2008. Field strength dependence of R1 and R relaxivities of human whole blood to prohance, vasovist, and deoxyhemoglobin. Magnetic Resonance in Medicine. 60, 1313–1320. https://doi.org/10.1002/mrm.21792.

Boas, D.A., Jones, S.R., Devor, A., Huppert, T.J., Dale, A.M., 2008. A vascular anatomical network model of the spatio-temporal response to brain activation. Neuroimage. 40, 1116–1129. https://doi.org/10.1016/j.neuroimage.2007.12.061.

Bollmann, S., Barth, M., 2020. New acquisition techniques and their prospects for the achievable resolution of fMRI. Progress in Neurobiology., 101936. https://doi.org/10.1016/j.pneurobio.2020.101936.

Buxton, R.B., 2009. Introduction to functional magnetic resonance imaging: principles and techniques. Cambridge university press.

Buxton, R.B., Uludağ, K., Dubowitz, D.J., Liu, T.T., 2004. Modeling the hemodynamic response to brain activation. Neuroimage. 23, S220–S233. https://doi.org/10.1016/j.neuroimage.2004.07.013.

Buxton, R.B., Wong, E.C., Frank, L.R., 1998. Dynamics of blood flow and oxygenation changes during brain activation: the balloon model. Magnetic Resonance in Medicine. 39, 855–864. https://doi.org/10.1002/mrm.1910390602.

Cassot, F., Lauwers, F., Lorthios, S., Puwanarajah, P., Duvernoy, H., 2009. Scaling laws for branching vessels of human cerebral cortex. Microcirculation. 16, 331–344. https://doi.org/10.1080/10739680802662607.

Chai, Y., Li, L., Huber, L., Poser, B.A., Bandettini, P.A., 2019. Integrated VASO and perfusion contrast: A new tool for laminar functional MRI. Neuroimage., 116358. https://doi.org/10.1016/j.neuroimage.2019.116358.

Chen, G., Wang, F., Gore, J.C., Roe, A.W., 2013. Layer-specific BOLD activation in awake monkey V1 revealed by ultra-high spatial resolution functional magnetic resonance imaging. Neuroimage. 64, 147–155. https://doi.org/10.1016/j.neuroimage.2012.08.060.

Chen, J.J., Pike, G.B., 2009. BOLD-specific cerebral blood volume and blood flow changes during neuronal activation in humans. NMR in Biomedicine 22, 1054–1062. https://doi.org/10.1002/nbm.1411.

Chen, J.J., Pike, G.B., 2010. MRI measurement of the BOLD-specific flow–volume relationship during hypercapnia and hypocapnia in humans. Neuroimage 53, 383–391. https://doi.org/10.1016/j.neuroimage.2010.07.003.

de Hollander, G., van der Zwaag, W., Qian, C., Zhang, P., Knapen, T., 2021. Ultra-high field fMRI reveals origins of feedforward and feedback activity within laminae of human ocular dominance columns. Neuroimage. 228, 117683. https://doi.org/10.1016/j.neuroimage.2020.117683.

De Martino, F., Zimmermann, J., Muckli, L., Ugurbil, K., Yacoub, E., Goebel, R., 2013. Cortical depth dependent functional responses in humans at 7T: improved specificity with 3D GRASE. PloS one 8, e60514. https://doi.org/10.1371/journal.pone.0060514.

Dobre, M.C., Uğurbil, K., Marjanska, M., 2007. Determination of blood longitudinal relaxation time (T1) at high magnetic field strengths. Magnetic Resonance Imaging. 25, 733–735. https://doi.org/10.1016/j.mri.2006.10.020.

Douglas, R.J., Martin, K.A., 2004. Neuronal circuits of the neocortex. Annu. Rev. Neurosci. 27, 419–451. https://doi.org/10.1146/annurev.neuro.27.070203.144152.

Duann, J.-R., Jung, T.-P., Kuo, W.-J., Yeh, T.-C., Makeig, S., Hsieh, J.-C., Sejnowski, T.J., 2002. Single-trial variability in event-related BOLD signals. Neuroimage. 15, 823–835. https://doi.org/10.1006/nimg.2001.1049.

Duvernoy, H.M., Delon, S., Vannson, J., 1981. Cortical blood vessels of the human brain. Brain Research Bulletin. 7, 519–579. https://doi.org/10.1016/0361-9230(81)90007-1.

Finn, E.S., Huber, L., Jangraw, D.C., Molfese, P.J., Bandettini, P.A., 2019. Layer-dependent activity in human prefrontal cortex during working memory. Nature Neuroscience. 22, 1687–1695. https://doi.org/10.1038/s41593-019-0487-z.

Fischl, B., Dale, A.M., 2000. Measuring the thickness of the human cerebral cortex from magnetic resonance images. Proceedings of the National Academy of Sciences. 97, 11050–11055. https://doi.org/10.1073/pnas.200033797.

Fracasso, A., Luijten, P.R., Dumoulin, S.O., Petridou, N., 2018. Laminar imaging of positive and negative BOLD in human visual cortex at 7 T. Neuroimage. 164, 100–111. https://doi.org/10.1016/j.neuroimage.2017.02.038.

Gagnon, L., Sakadžić, S., Lesage, F., Musacchia, J.J., Lefebvre, J., Fang, Q., Yücel, M.A., Evans, K.C., Mandeville, E.T., Cohen-Adad, J., 2015. Quantifying the microvascular origin of BOLD-fMRI from first principles with two-photon microscopy and an oxygen-sensitive nanoprobe. Journal of Neuroscience. 35, 3663–3675. https://doi.org/10.1523/JNEUROSCI.3555-14.2015.

Genois, É., Gagnon, L., Desjardins, M., 2020. Modeling of vascular space occupancy and BOLD functional MRI from first principles using real microvascular angiograms. Magnetic Resonance in Medicine. 85, 456–468. https://doi.org/10.1002/mrm.28429.

Goense, J., Merkle, H., Logothetis, N.K., 2012. High-resolution fMRI reveals laminar differences in neurovascular coupling between positive and negative BOLD responses. Neuron. 76, 629–639. https://doi.org/10.1016/j.neuron.2012.09.019.

Goense, J.B., Logothetis, N.K., 2006. Laminar specificity in monkey V1 using high-resolution SE-fMRI. Magnetic Resonance Imaging. 24, 381–392. https://doi.org/10.1016/j.mri.2005.12.032.

Griffeth, V.E., Simon, A.B., Buxton, R.B., 2015. The coupling of cerebral blood flow and oxygen metabolism with brain activation is similar for simple and complex stimuli in human primary visual cortex. Neuroimage. 104, 156–162. https://doi.org/10.1016/j.neuroimage.2014.10.003.

Griswold, M.A., Jakob, P.M., Heidemann, R.M., Nittka, M., Jellus, V., Wang, J., Kiefer, B., Haase, A., 2002. Generalized autocalibrating partially parallel acquisitions (GRAPPA). Magnetic Resonance in Medicine. 47, 1202–1210. https://doi.org/10.1002/mrm.10171.

Grubb, R.L., Raichle, M.E., Eichling, J.O., Ter-Pogossian, M.M., 1974. The effects of changes in PaCO2 cerebral blood volume, blood flow, and vascular mean transit time. Stroke. 5, 630–639. https://doi.org/10.1161/01.STR.5.5.630.

Handwerker, D.A., Ollinger, J.M., D’Esposito, M., 2004. Variation of BOLD hemodynamic responses across subjects and brain regions and their effects on statistical analyses. Neuroimage. 21, 1639–1651. https://doi.org/10.1016/j.neuroimage.2003.11.029.

Havlicek, M., Uludağ, K., 2020. A dynamical model of the laminar BOLD response. Neuroimage. 204, 116209. https://doi.org/10.1016/j.neuroimage.2019.116209.

Heinzle, J., Koopmans, P.J., den Ouden, H.E., Raman, S., Stephan, K.E., 2016. A hemodynamic model for layered BOLD signals. Neuroimage. 125, 556–570. https://doi.org/10.1016/j.neuroimage.2015.10.025.

Hillman, E.M., Devor, A., Bouchard, M.B., Dunn, A.K., Krauss, G., Skoch, J., Bacskai, B.J., Dale, A.M., Boas, D.A., 2007. Depth-resolved optical imaging and microscopy of vascular compartment dynamics during somatosensory stimulation. Neuroimage. 35, 89–104. https://doi.org/10.1016/j.neuroimage.2006.11.032.

Huber, L., Goense, J., Ivanov, D., Krieger, S., Turner, R., Moeller, H.E., 2013. Cerebral blood volume changes in negative BOLD regions during visual stimulation in humans at 7T. 21st Annual Meeting of the International Society for Magnetic Resonance in Medicine.

Huber, L., Goense, J., Kennerley, A.J., Ivanov, D., Krieger, S.N., Lepsien, J., Trampel, R., Turner, R., Möller, H.E., 2014a. Investigation of the neurovascular coupling in positive and negative BOLD responses in human brain at 7 T. Neuroimage. 97, 349–362. https://doi.org/10.1016/j.neuroimage.2014.04.022.

Huber, L., Goense, J., Kennerley, A.J., Trampel, R., Guidi, M., Reimer, E., Ivanov, D., Neef, N., Gauthier, C.J., Turner, R., 2015. Cortical lamina-dependent blood volume changes in human brain at 7 T. Neuroimage. 107, 23–33. https://doi.org/10.1016/j.neuroimage.2014.11.046.

Huber, L., Handwerker, D.A., Jangraw, D.C., Chen, G., Hall, A., Stüber, C., Gonzalez-Castillo, J., Ivanov, D., Marrett, S., Guidi, M., 2017a. High-resolution CBV-fMRI allows mapping of laminar activity and connectivity of cortical input and output in human M1. Neuron. 96, 1253-1263. e1257. https://doi.org/10.1016/j.neuron.2017.11.005.

Huber, L., Ivanov, D., Handwerker, D.A., Marrett, S., Guidi, M., Uludağ, K., Bandettini, P.A., Poser, B.A., 2016. Techniques for blood volume fMRI with VASO: from low-resolution mapping towards sub-millimeter layer-dependent applications. Neuroimage. https://doi.org/10.1016/j.neuroimage.2016.11.039.

Huber, L., Ivanov, D., Krieger, S.N., Streicher, M.N., Mildner, T., Poser, B.A., Möller, H.E., Turner, R., 2014b. Slab-selective, BOLD-corrected VASO at 7 Tesla provides measures of cerebral blood volume reactivity with high signal-to-noise ratio. Magnetic Resonance in Medicine. 72, 137–148. https://doi.org/10.1002/mrm.24916.

Huber, L., Uludağ, K., Möller, H.E., 2017b. Non-BOLD contrast for laminar fMRI in humans: CBF, CBV, and CMRO2. Neuroimage. https://doi.org/10.1016/j.neuroimage.2017.07.041.

Huber, L.R., Poser, B.A., Bandettini, P.A., Arora, K., Wagstyl, K., Cho, S., Goense, J., Nothnagel, N., Morgan, A.T., van den Hurk, J., 2021. LAYNII: a software suite for layer-fMRI. Neuroimage 237, 118091. https://doi.org/10.1016/j.neuroimage.2021.118091.

Ito, H., Ibaraki, M., Kanno, I., Fukuda, H., Miura, S., 2005. Changes in the arterial fraction of human cerebral blood volume during hypercapnia and hypocapnia measured by positron emission tomography. Journal of Cerebral Blood Flow & Metabolism. 25, 852–857. https://doi.org/10.1038%2Fsj.jcbfm.9600076.

Ito, H., Kanno, I., Iida, H., Hatazawa, J., Shimosegawa, E., Tamura, H., Okudera, T., 2001. Arterial fraction of cerebral blood volume in humans measured by positron emission tomography. Annals of Nuclear Medicine. 15, 111–116. https://doi.org/10.1007/BF02988600.

Jin, T., Kim, S.-G., 2006. Spatial dependence of CBV-fMRI: a comparison between VASO and contrast agent based methods. 2006 International Conference of the IEEE Engineering in Medicine and Biology Society. IEEE, pp. 25–28. https://doi.org/10.1109/IEMBS.2006.259553.

Jin, T., Kim, S.-G., 2008a. Cortical layer-dependent dynamic blood oxygenation, cerebral blood flow and cerebral blood volume responses during visual stimulation. Neuroimage. 43, 1–9. https://doi.org/10.1016/j.neuroimage.2008.06.029.

Jin, T., Kim, S.-G., 2008b. Improved cortical-layer specificity of vascular space occupancy fMRI with slab inversion relative to spin-echo BOLD at 9.4 T. Neuroimage. 40, 59–67. https://doi.org/10.1016/j.neuroimage.2007.11.045.

Kashyap, S., Ivanov, D., Havlicek, M., Poser, B.A., Uludağ, K., 2018. Impact of acquisition and analysis strategies on cortical depth-dependent fMRI. Neuroimage 168, 332–344. https://doi.org/10.1016/j.neuroimage.2018.02.027.

Kim, S.G., Hendrich, K., Hu, X., Merkle, H., Ugurbil, K., 1994. Potential pitfalls of functional MRI using conventional gradient-recalled echo techniques. NMR in Biomedicine. 7, 69–74. https://doi.org/10.1002/nbm.1940070111.

Kim, T., Kim, S.-G., 2010. Cortical layer-dependent arterial blood volume changes: improved spatial specificity relative to BOLD fMRI. Neuroimage. 49, 1340–1349. https://doi.org/10.1016/j.neuroimage.2009.09.061.

Kim, T., Kim, S.-G., 2011a. Suppl 1: Quantitative MRI of Cerebral Arterial Blood Volume. The Open Neuroimaging Journal. 5, 136. https://dx.doi.org/10.2174%2F1874440001105010136.

Kim, T., Kim, S.-G., 2011b. Temporal dynamics and spatial specificity of arterial and venous blood volume changes during visual stimulation: implication for BOLD quantification. Journal of Cerebral Blood Flow & Metabolism. 31, 1211–1222. https://doi.org/10.1038%2Fjcbfm.2010.226.

Klein, B.P., Fracasso, A., van Dijk, J.A., Paffen, C.L., Te Pas, S.F., Dumoulin, S.O., 2018. Cortical depth dependent population receptive field attraction by spatial attention in human V1. Neuroimage 176, 301–312. https://doi.org/10.1016/j.neuroimage.2018.04.055.

Koopmans, P.J., Barth, M., Norris, D.G., 2010. Layer-specific BOLD activation in human V1. Human Brain Mapping. 31, 1297–1304. https://doi.org/10.1002/hbm.20936.

Koopmans, P.J., Barth, M., Orzada, S., Norris, D.G., 2011. Multi-echo fMRI of the cortical laminae in humans at 7 T. Neuroimage. 56, 1276–1285. https://doi.org/10.1016/j.neuroimage.2011.02.042.

Lauwers, F., Cassot, F., Lauwers-Cances, V., Puwanarajah, P., Duvernoy, H., 2008. Morphometry of the human cerebral cortex microcirculation: general characteristics and space-related profiles. Neuroimage. 39, 936–948. https://doi.org/10.1016/j.neuroimage.2007.09.024.

Lawrence, S.J., Norris, D.G., De Lange, F.P., 2019. Dissociable laminar profiles of concurrent bottom-up and top-down modulation in the human visual cortex. Elife. 8, e44422. https://doi.org/10.7554/eLife.44422.

Light, K.C., Turner, J.R., Hinderliter, A.L., Sherwood, A., 1993. Race and gender comparisons: I. Hemodynamic responses to a series of stressors. Health Psychology. 12, 354. https://psycnet.apa.org/doi/10.1037/0278-6133.12.5.354.

Lu, H., Golay, X., Pekar, J.J., van Zijl, P., 2003. Functional magnetic resonance imaging based on changes in vascular space occupancy. Magnetic Resonance in Medicine. 50, 263–274. https://doi.org/10.1002/mrm.10519.

Lu, H., Hua, J., Zijl, P., 2013. Noninvasive functional imaging of cerebral blood volume with vascular-space-occupancy (VASO) MRI. NMR in Biomedicine. 26, 932–948. https://doi.org/10.1002/nbm.2905.

Mandeville, J.B., Marota, J.J., Ayata, C., Zaharchuk, G., Moskowitz, M.A., Rosen, B.R., Weisskoff, R.M., 1999. Evidence of a cerebrovascular postarteriole windkessel with delayed compliance. Journal of Cerebral Blood Flow & Metabolism. 19, 679–689. https://doi.org/10.1097%2F00004647-199906000-00012.

Markuerkiaga, I., Barth, M., Norris, D.G., 2016. A cortical vascular model for examining the specificity of the laminar BOLD signal. Neuroimage. 132, 491–498. https://doi.org/10.1016/j.neuroimage.2016.02.073.

Markuerkiaga, I., Marques, J.P., Gallagher, T.E., Norris, D.G., 2021. Estimation of laminar BOLD activation profiles using deconvolution with a physiological point spread function. Journal of Neuroscience Methods. 353, 109095. https://doi.org/10.1016/j.jneumeth.2021.109095.

Marquardt, I., De Weerd, P., Schneider, M., Gulban, O.F., Ivanov, D., Wang, Y., Uludağ, K., 2020. Feedback contribution to surface motion perception in the human early visual cortex. Elife 9, e50933. https://doi.org/10.7554/eLife.50933.

Marques, J.P., Kober, T., Krueger, G., van der Zwaag, W., Van de Moortele, P.-F., Gruetter, R., 2010. MP2RAGE, a self bias-field corrected sequence for improved segmentation and T1-mapping at high field. Neuroimage. 49, 1271–1281. https://doi.org/10.1016/j.neuroimage.2009.10.002.

Norris, D.G., Polimeni, J.R., 2019. Laminar (f) MRI: A short history and future prospects. Neuroimage. https://doi.org/10.1016/j.neuroimage.2019.04.082.

O’Brien, K.R., Magill, A.W., Delacoste, J., Marques, J.P., Kober, T., Fautz, H.P., Lazeyras, F., Krueger, G., 2014. Dielectric pads and low-adiabatic pulses: Complementary techniques to optimize structural T1w whole-brain MP2RAGE scans at 7 tesla. Magnetic Resonance Imaging. 40, 804–812. https://doi.org/10.1002/jmri.24435.

Obata, T., Liu, T.T., Miller, K.L., Luh, W.-M., Wong, E.C., Frank, L.R., Buxton, R.B., 2004. Discrepancies between BOLD and flow dynamics in primary and supplementary motor areas: application of the balloon model to the interpretation of BOLD transients. Neuroimage. 21, 144–153. https://doi.org/10.1016/j.neuroimage.2003.08.040.

Ogawa, S., Lee, T.-M., Kay, A.R., Tank, D.W., 1990. Brain magnetic resonance imaging with contrast dependent on blood oxygenation. Proceedings of the National Academy of Sciences. 87, 9868–9872. https://doi.org/10.1073/pnas.87.24.9868.

Oliveira, Í.A., van der Zwaag, W., Raimondo, L., Dumoulin, S.O., Siero, J.C., 2021a. Comparing hand movement rate dependence of cerebral blood volume and BOLD responses at 7T. Neuroimage 226, 117623.

Oliveira, Í.A., van der Zwaag, W., Raimondo, L., Dumoulin, S.O., Siero, J.C., 2021b. Comparing hand movement rate dependence of cerebral blood volume and BOLD responses at 7T. Neuroimage. 226, 117623. https://doi.org/10.1016/j.neuroimage.2020.117623.

Olman, C.A., Harel, N., Feinberg, D.A., He, S., Zhang, P., Ugurbil, K., Yacoub, E., 2012. Layer-specific fMRI reflects different neuronal computations at different depths in human V1. PloS one. 7, e32536. https://doi.org/10.1371/journal.pone.0032536.

Petridou, N., Siero, J.C., 2017. Laminar fMRI: What can the time domain tell us? Neuroimage. https://doi.org/10.1016/j.neuroimage.2017.07.040.

Polimeni, J.R., Fischl, B., Greve, D.N., Wald, L.L., 2010. Laminar analysis of 7 T BOLD using an imposed spatial activation pattern in human V1. Neuroimage. 52, 1334–1346. https://doi.org/10.1016/j.neuroimage.2010.05.005.

Polimeni, J.R., Hinds, O.P., Balasubramanian, M., van der Kouwe, A., Wald, L.L., Dale, A.M., Fischl, B., Schwartz, E.L., 2005. The human V1–V2–V3 visuotopic map complex measured via fMRI at 3 and 7 Tesla.

Polimeni, J.R., Renvall, V., Zaretskaya, N., Fischl, B., 2018. Analysis strategies for high-resolution UHF-fMRI data. Neuroimage. 168, 296–320. https://doi.org/10.1016/j.neuroimage.2017.04.053.

Poplawsky, A.J., Fukuda, M., Murphy, M., Kim, S.-G., 2015. Layer-specific fMRI responses to excitatory and inhibitory neuronal activities in the olfactory bulb. Journal of Neuroscience. 35, 15263–15275. https://doi.org/10.1523/JNEUROSCI.1015-15.2015.

Poplawsky, A.J., Kim, S.-G., 2014. Layer-dependent BOLD and CBV-weighted fMRI responses in the rat olfactory bulb. Neuroimage. 91, 237–251. https://doi.org/10.1016/j.neuroimage.2013.12.067.

Poser, B.A., Koopmans, P.J., Witzel, T., Wald, L.L., Barth, M., 2010. Three dimensional echo-planar imaging at 7 Tesla. Neuroimage. 51, 261–266. https://doi.org/10.1016/j.neuroimage.2010.01.108.

Ress, D., Glover, G.H., Liu, J., Wandell, B., 2007. Laminar profiles of functional activity in the human brain. Neuroimage. 34, 74–84. https://doi.org/10.1016/j.neuroimage.2006.08.020.

Schmid, F., Barrett, M.J., Jenny, P., Weber, B., 2019. Vascular density and distribution in neocortex. Neuroimage. https://doi.org/10.1016/j.neuroimage.2017.06.046.

Self, M.W., van Kerkoerle, T., Goebel, R., Roelfsema, P.R., 2017. Benchmarking laminar fMRI: neuronal spiking and synaptic activity during top-down and bottom-up processing in the different layers of cortex. Neuroimage. https://doi.org/10.1016/j.neuroimage.2017.06.045.

Shih, Y.-Y.I., Chen, Y.-Y., Lai, H.-Y., Kao, Y.-C.J., Shyu, B.-C., Duong, T.Q., 2013. Ultra high-resolution fMRI and electrophysiology of the rat primary somatosensory cortex. Neuroimage. 73, 113–120. https://doi.org/10.1016/j.neuroimage.2013.01.062.

Silva, A.C., Koretsky, A.P., 2002. Laminar specificity of functional MRI onset times during somatosensory stimulation in rat. Proceedings of the National Academy of Sciences. 99, 15182–15187. https://doi.org/10.1073/pnas.222561899.

Silva, A.C., Koretsky, A.P., Duyn, J.H., 2007. Functional MRI impulse response for BOLD and CBV contrast in rat somatosensory cortex. Magnetic Resonance in Medicine. 57, 1110–1118. https://doi.org/10.1002/mrm.21246.

Stefanovic, B., Pike, G.B., 2005. Venous refocusing for volume estimation: VERVE functional magnetic resonance imaging. Magnetic Mesonance in Medicine. 53, 339–347. https://doi.org/10.1002/mrm.20352.

Stephan, K.E., Petzschner, F.H., Kasper, L., Bayer, J., Wellstein, K.V., Stefanics, G., Pruessmann, K.P., Heinzle, J., 2019. Laminar fMRI and computational theories of brain function. Neuroimage. 197, 699–706. https://doi.org/10.1016/j.neuroimage.2017.11.001.

Takano, T., Tian, G.-F., Peng, W., Lou, N., Libionka, W., Han, X., Nedergaard, M., 2006. Astrocyte-mediated control of cerebral blood flow. Nature Neuroscience. 9, 260. https://doi.org/10.1038/nn1623.

Turner, R., 2002. How much cortex can a vein drain? Downstream dilution of activation-related cerebral blood oxygenation changes. Neuroimage. 16, 1062–1067. https://doi.org/10.1006/nimg.2002.1082.

Uludağ, K., Müller-Bierl, B., Uğurbil, K., 2009. An integrative model for neuronal activity-induced signal changes for gradient and spin echo functional imaging. Neuroimage. 48, 150–165. https://doi.org/10.1016/j.neuroimage.2009.05.051.

van Dijk, J.A., Fracasso, A., Petridou, N., Dumoulin, S.O., 2020. Linear systems analysis for laminar fMRI: Evaluating BOLD amplitude scaling for luminance contrast manipulations. Scientific Reports. 10, 1–15. https://doi.org/10.1038/s41598-020-62165-x.

Van Kerkoerle, T., Self, M.W., Roelfsema, P.R.J.N.c., 2017. Layer-specificity in the effects of attention and working memory on activity in primary visual cortex. Nature Communications. 8, 1–14. 10.1038/ncomms13804.

Vanzetta, I., Hildesheim, R., Grinvald, A., 2005. Compartment-resolved imaging of activity-dependent dynamics of cortical blood volume and oximetry. Journal of Neuroscience. 25, 2233–2244. https://doi.org/10.1523/JNEUROSCI.3032-04.2005.

Vizioli, L., De Martino, F., Petro, L.S., Kersten, D., Ugurbil, K., Yacoub, E., Muckli, L., 2020. Multivoxel Pattern of Blood Oxygen Level Dependent Activity can be sensitive to stimulus specific fine scale responses. Scientific Reports. 10, 1–18. https://doi.org/10.1038/s41598-020-64044-x.

Waehnert, M., Dinse, J., Weiss, M., Streicher, M.N., Waehnert, P., Geyer, S., Turner, R., Bazin, P.-L., 2014. Anatomically motivated modeling of cortical laminae. Neuroimage. 93, 210–220. https://doi.org/10.1016/j.neuroimage.2013.03.078.

Weber, B., Keller, A.L., Reichold, J., Logothetis, N.K., 2008. The microvascular system of the striate and extrastriate visual cortex of the macaque. Cerebral Cortex. 18, 2318–2330. https://doi.org/10.1093/cercor/bhm259.

Yablonskiy, D.A., Haacke, E.M., 1994. Theory of NMR signal behavior in magnetically inhomogeneous tissues: the static dephasing regime. Magnetic Resonance in Medicine. 32, 749–763. https://doi.org/10.1002/mrm.1910320610.

Yu, X., Qian, C., Chen, D.-y., Dodd, S.J., Koretsky, A.P., 2014. Deciphering laminar-specific neural inputs with line-scanning fMRI. Nature Methods. 11, 55. https://doi.org/10.1038/nmeth.2730.

Zaretskaya, N., Bause, J., Polimeni, J.R., Grassi, P.R., Scheffler, K., Bartels, A., 2020. Eye-selective fMRI activity in human primary visual cortex: Comparison between 3 T and 9.4 T, and effects across cortical depth. Neuroimage. 220, 117078. https://doi.org/10.1016/j.neuroimage.2020.117078.

Zhang, X., Petersen, E.T., Ghariq, E., De Vis, J., Webb, A., Teeuwisse, W.M., Hendrikse, J., Van Osch, M., 2013. In vivo blood T1 measurements at 1.5 T, 3 T, and 7 T. Magnetic Resonance in Medicine 70, 1082–1086. https://doi.org/10.1002/mrm.24550.

Zhao, F., Wang, P., Hendrich, K., Ugurbil, K., Kim, S.-G., 2006. Cortical layer-dependent BOLD and CBV responses measured by spin-echo and gradient-echo fMRI: insights into hemodynamic regulation. Neuroimage. 30, 1149–1160. https://doi.org/10.1016/j.neuroimage.2005.11.013.

Zweifach, B.W., Lipowsky, H.H., 1977. Quantitative studies of microcirculatory structure and function. III. Microvascular hemodynamics of cat mesentery and rabbit omentum. Circulation Research. 41, 380–390. https://doi.org/10.1161/01.RES.41.3.380.

