## supplementary Materials for "Modelling the depth-dependent VASO and BOLD responses in human primary visual cortex"

### 1 Appendix A. Supplementary Materials

#### 1.1 The Effect of the Constant and Depth-Dependent Baseline CBV on the Simulated Profiles

We also simulated the VASO and BOLD responses with constant and depth-dependent baseline CBV in the laminar network to show the effect of the baseline CBV on the simulated profiles. In addition, we also assumed two patterns of “equal activation strength across the layers” and “higher activation strength in the middle cortical depth” in the laminar network. The four simulated scenarios are: (A) depth-dependent baseline CBV in the laminar network and higher activation strength in middle depths, (B) depth-dependent baseline CBV of the laminar network and equal activation strength across all depths, (C) Constant baseline CBV of the laminar network and higher activation strength in middle depths, (D) Constant baseline CBV of the laminar network and equal activation strength across all depths. These simulations were performed using the CBV and oxygenation changes reported in Table 3. For more detail about the simulation assumptions see section 2.3 and 2.4.

In detail, scenario (A) assumes a depth-dependent baseline CBV in the laminar network (Figure 1B), an increased CBV change in the middle layers (Figure 2), and ΔCBV and oxygenation changes as reported in Table 4. Further, scenario (B) also assumes a depth-dependent baseline CBV in the laminar network (Figure 1B), but equal CBV change across all layers (Figure S2), and ΔCBV and oxygenation changes as reported in Table 4. In contrast, scenarios (C) and (D) assume a constant baseline CBV in the laminar network (Figure 3). Moreover, scenario (C) assumes an increased CBV change in the middle layers (Figure 2), whereas scenario (D) assumes equal CBV change across all layers (Figure S2). Both use the ΔCBV and oxygenation changes as reported in Table 4.

Figure S1 demonstrates the results of these simulations with the lowest RMSE (best fit), along with the measured profiles. Scenarios of (B) and (D) failed to explain the expected VASO profiles from the imaging data, showing the sensitivity of the VASO response to the $\Delta CBV$ across the layers, while it is relatively insensitive to the baseline CBV (be it constant or depth-dependent). The so-called “bump” in the simulated BOLD response is more prominent in scenarios of (A) and (B) – where baseline CBV is a function of the cortical depth – compared to scenarios of (C) and (D), showing the increased influence of baseline CBV on the simulated BOLD response.

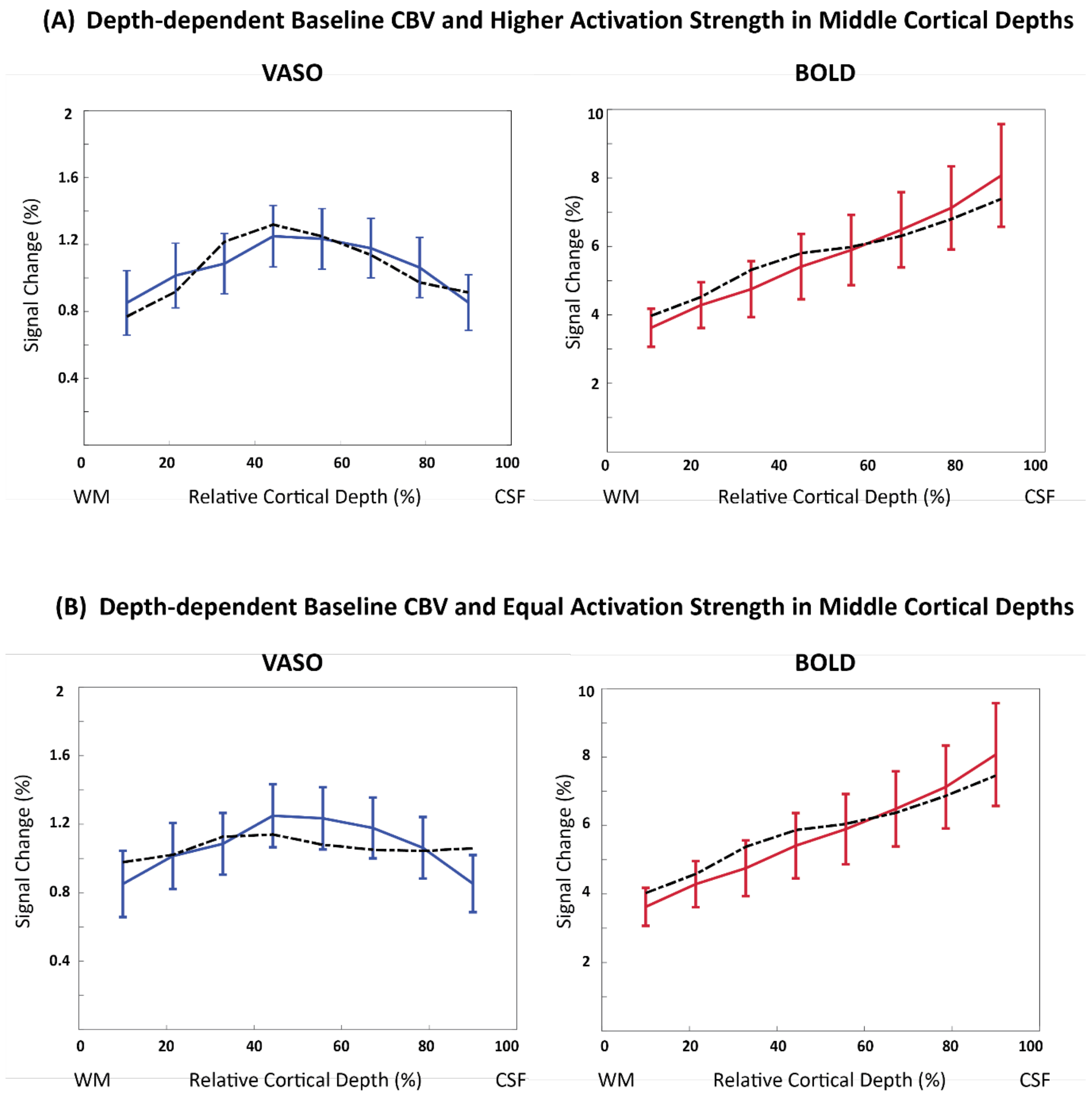

**(C) Constant Baseline CBV and Higher Activation Strength in Middle Cortical Depths**

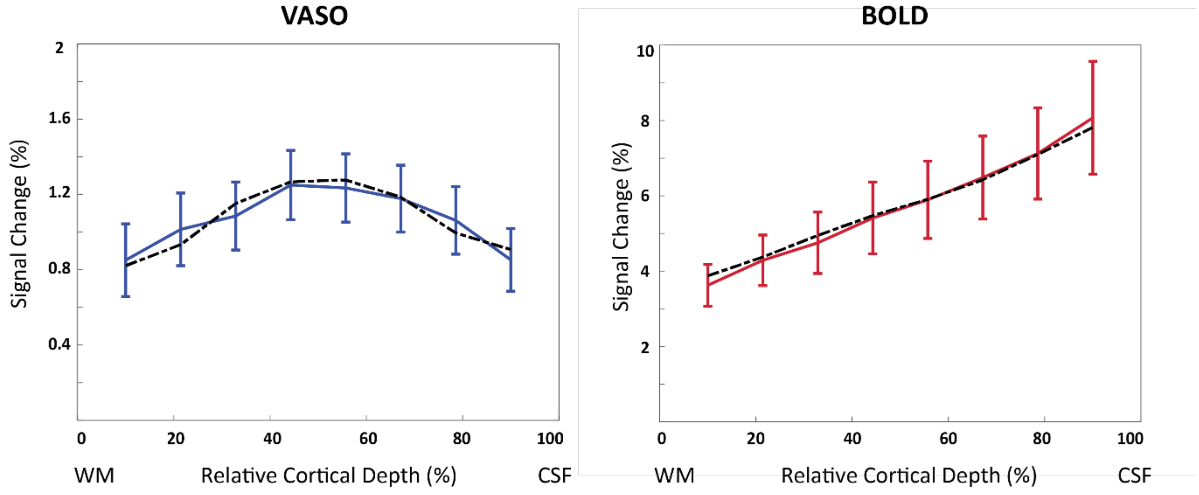


**(D) Constant Baseline CBV and Equal Activation Strength in Middle Cortical Depths**

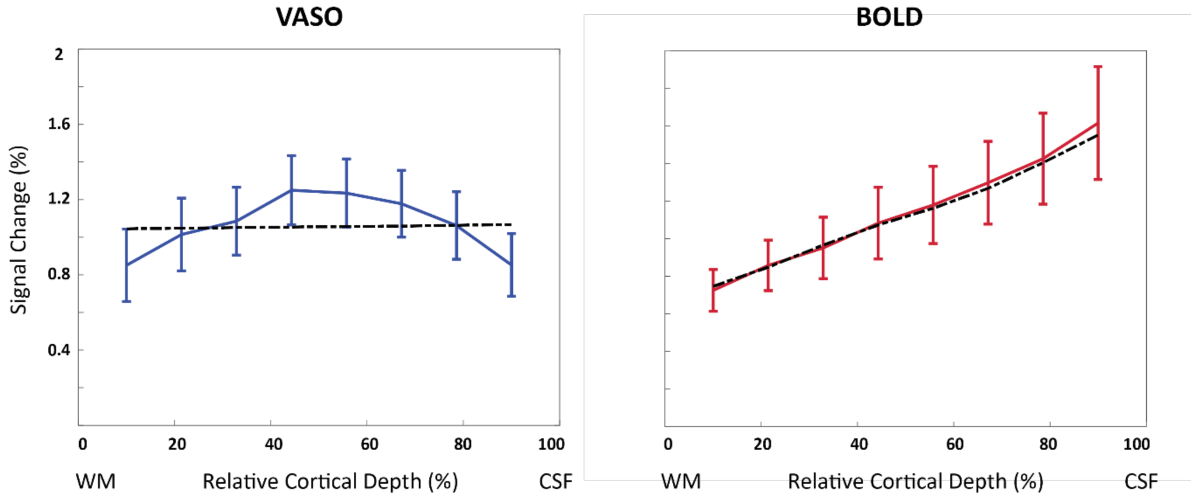


Figure S1: The simulated VASO and BOLD responses in four scenarios of: (A) depth- dependent baseline CBV in the laminar network and higher activation strength in middle depths. (B) depth-dependent baseline CBV of the laminar network and equal activation strength across the layers. (C) Constant baseline CBV (2.3%) of the laminar network and higher activation strength in middle depths. (D) Constant baseline CBV (2.3%) of the laminar network and equal activation strength in middle depths.

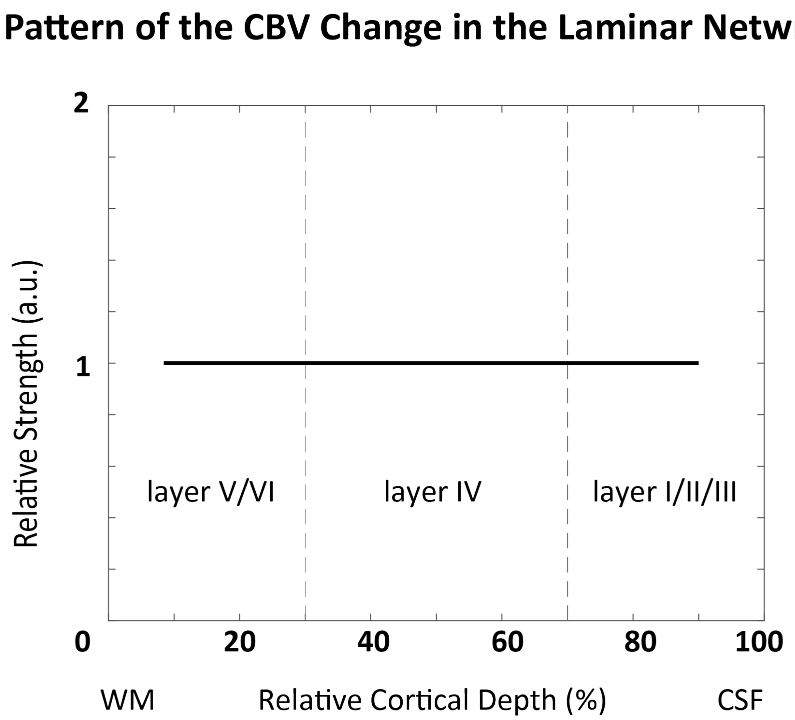


Figure S2: The pattern of the CBV change in the laminar network used to simulate the VASO and BOLD responses in scenarios of B and D in section 1.1 of Appendix A.

#### 1.2 RMSE Plots Corresponding to the Scenario Shown in Figure 2

To investigate the sensitivity of the model to the choice of input parameters, we plotted the profiles with RMSEs that are 20% higher than the minimum RMSE as the shaded area in Figure 6. The RMSE values of simulated VASO and BOLD profiles for different input parameter combinations are shown in Figure S3. To make the comparison easier across different scenarios, we have presented the percentage RMSEs normalized by the minimum RMSE in each scenario. Note that the input parameters for VASO simulations are $\Delta CBV$ in ICAs, arterioles/capillaries, venules, and ICVs, and the input for the BOLD simulation are blood oxygenation values ($Y$) during baseline and activation in venules and ICVs. The white dotted lines indicate the range of RMSE values up to 20% higher than the minimum RMSE. Note how a wide diagonal band of minimal RMSE values in Figure S3A is contained within this limit, which indicates that a large number of possible $\Delta CBV$ combinations would result in similar profiles, ranging from no change to 90% CBV change in venules, and 40% to 60% $\Delta CBV$ in arterioles and capillaries. Smaller regions of minimal RMSE can be found in Figure S3B-E, indicating that a narrower range of $\Delta CBV$ combinations would give rise to similar profiles for these parameter combinations. Again, in Figure S3F, a wide diagonal band indicates a large number of combinations of $\Delta CBV$ values in ICAs and ICVs which would result in similar VASO profiles. This shows a high differentiability between macro- and micro-vascular compartments (Figure S3 B-F), but considerable interchangeability within each vascular class (Figures S3A and S3F).

The RMSE for the BOLD simulations with varying blood oxygenation values at baseline and during activation is shown in Figure S3G. Again, the diagonal band enclosed by the 20% RMSE limit indicates that various combinations of blood oxygenation values would result in similar profiles. A ΔY of approximately 20% between baseline and activation yields similarly low RMSE values.

#### VASO RMSEs Corresponding to the Figure 2

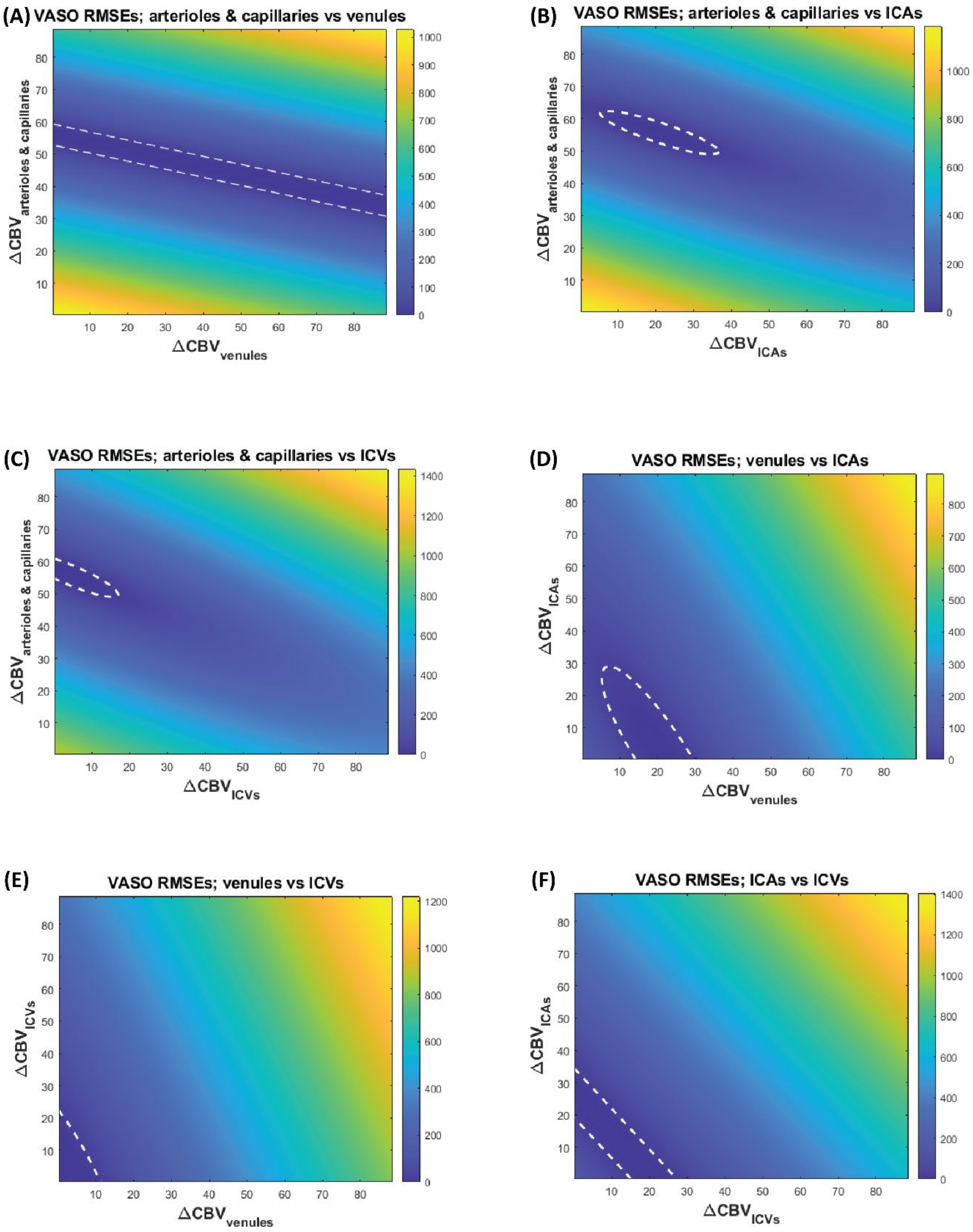

#### BOLD RMSEs Corresponding to the Figure 2

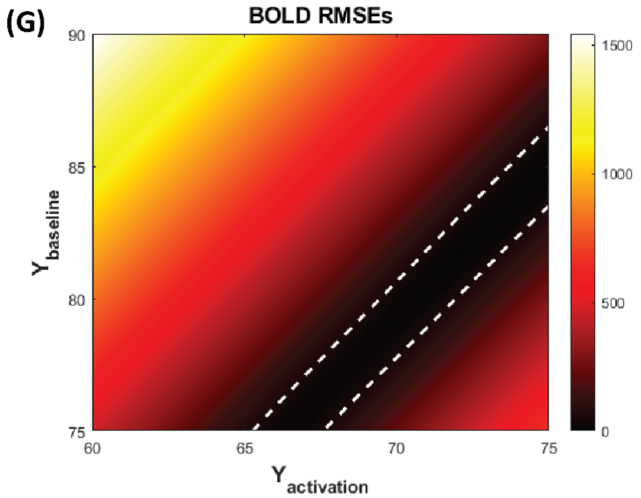


Figure S3: The root-mean-square-error (RMSE) values calculated as a measure of similarity between simulated and measured VASO (A-F) and BOLD (G) profiles correspond to Figure 6 of the manuscript. The white dotted lines refer to the RMSEs that are 20% higher than the minimum RMSE. (A) shows the RMSE when varying $\Delta CBV$ in arterioles/capillaries and venules but keeping the $\Delta CBV$ in other compartments at the value obtained from the profile with minimum RMSE. Similarly, (B) the $\Delta CBV$ in arterioles/capillaries and ICAs is varied but kept constant in ICVs and venules. (C) the RMSE is shown when varying the $\Delta CBV$ in arterioles/capillaries and ICVs but using the optimal value for the $\Delta CBV$ in ICAs and venules. (D) the RMSEs correspond to varying CBV change in venules and ICAs. (E) the RMSEs when varying the CBV change in venules and ICVs. (F) the RMSEs related to varying CBV change in ICAs and ICVs, while using the optimal value for the rest of the compartments. (G) BOLD RMSEs when varying the oxygen saturation at baseline and activation.

#### 1.3 RMSE Plots Corresponding to the Constant Baseline CBV Scenario

Figure S4 illustrates the VASO and BOLD RMSE plots corresponding to the constant baseline CBV scenario. The white dotted lines indicate the RMSEs that are 20% higher than the minimum RMSE. Note the smaller 20% regime compared to the scenario shown in Figure S3, particularly for the BOLD RMSE plot. Similar to Figure S3, differentiating *between* the micro-vessels in the laminar network (Figure S4 A) is challenging and results in a wide range of $\Delta CBV$ combinations that give rise to similar profiles for these parameter combinations. This 20% regime is much smaller in Figures S4 B-F, indicating the higher differentiability between the micro- and macro-vessels (i.e. the laminar network compartments vs ICAs/ICVs).

#### VASO RMSEs at the Constant Baseline CBV Scenario

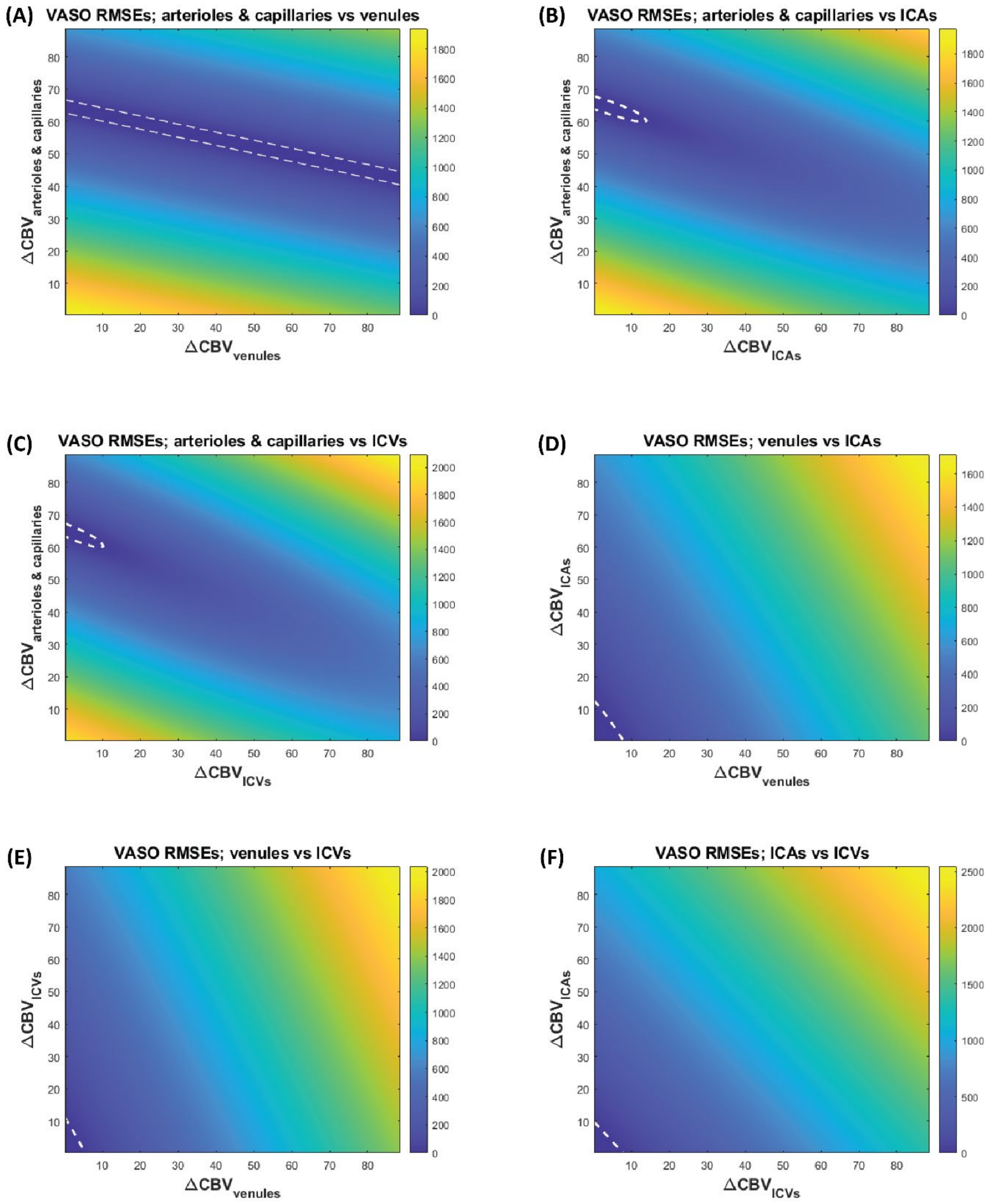

#### BOLD RMSEs at the Constant Baseline CBV Scenario

(G)

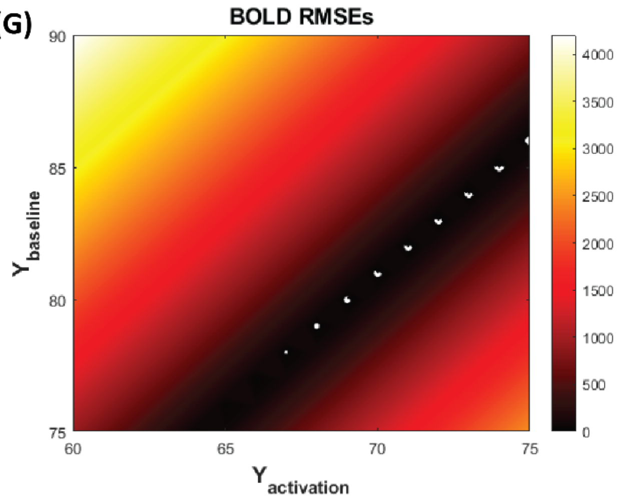


Figure S4: The root-mean-square-error (RMSE) values calculated as a measure of similarity between simulated and measured VASO (A-F) and BOLD (G) profiles correspond to the "constant baseline CBV" scenario. The white dotted lines refer to the RMSEs that are 20% higher than the minimum RMSE. (A) shows the RMSE when varying $\Delta CBV$ in arterioles/capillaries and venules but keeping the $\Delta CBV$ in other compartments at the value obtained from the profile with minimum RMSE. Similarly, (B) the $\Delta CBV$ in arterioles/capillaries and ICAs is varied but kept constant in ICVs and venules. (C) the RMSE is shown when varying the $\Delta CBV$ in arterioles/capillaries and ICVs but using the optimal value for the $\Delta CBV$ in ICAs and venules. (D) the RMSEs correspond to varying CBV change in venules and ICAs. (E) the RMSEs when varying the CBV change in venules and ICVs. (F) the RMSEs related to varying CBV change in ICAs and ICVs, while using the optimal value for the rest of the compartments. (G) BOLD RMSEs when varying the oxygen saturation at baseline and activation.

#### 1.4 Equal Activation Strength Across Depths in Intra-Cortical Vessels

Another possible scenario is equal activation strength in intra-cortical vessels (ICAs and ICVs) across cortical depth, while assuming a higher CBV change in the middle layers of the laminar network. Figure S5 shows the pattern of this activation strength in all vessel compartments and the corresponding simulated VASO and BOLD responses are shown in Figure S6. With this scenario, the VASO best-fit estimates 36% CBV change in arterioles and capillaries, 1% in venules, 12% in ICAs, and 14% in ICVs.

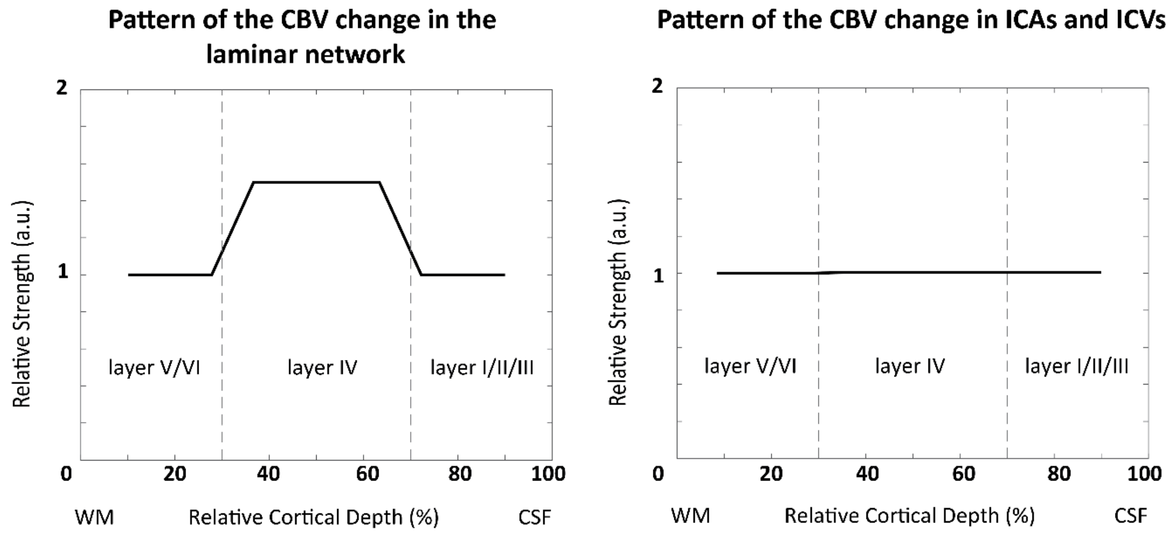


Figure S5: The pattern of the CBV change in the laminar network (left) and intra-cortical vessels (right) as one of the possible scenarios: higher CBV change in middle depths for the laminar network, and equal CBV change in ICAs and ICVs across all cortical layers.

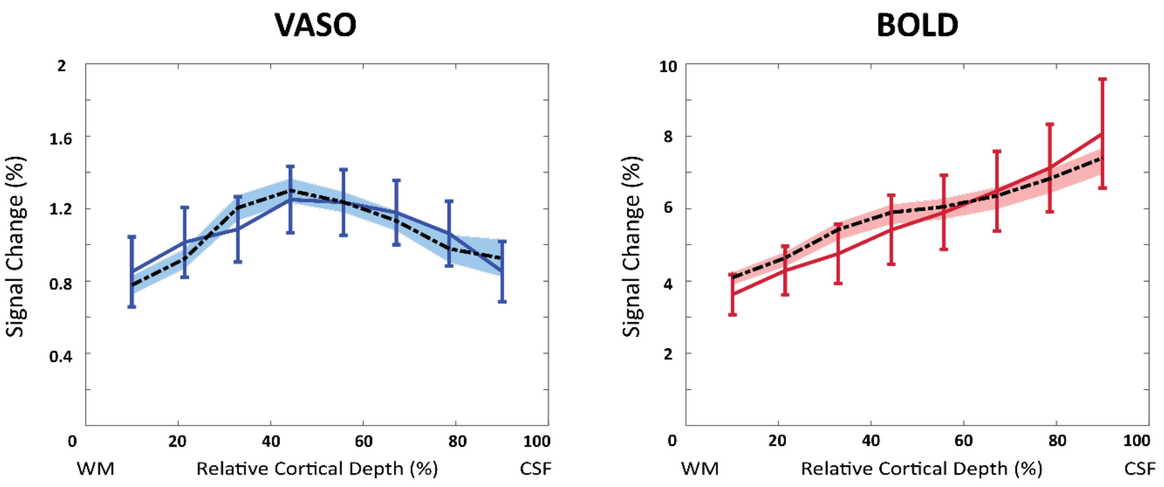


Figure S6: The VASO and BOLD best fit (dotted line), imaging data (solid line), and 20% regime (shaded area) correspond to scenario shown in Figure S4.

#### 1.5 Simulating the VASO Response with 250 μm Resolution and No Smoothing

To investigate the potential of the model to predict the double-peak VASO response obtained from macaque brain (Huber et al., 2015), we simulated the CBV response with 250μm resolution and no smoothing kernel assuming a depth-dependent baseline CBV (Figure 1), an increased CBV change in the middle layers (Figure 2), and ΔCBV and oxygenation changes as reported in Table 4. The two potential peaks correspond to higher local CBV change in both side of stria of Gennari that can possibly be observed with higher acquisition resolution is shown in Figure S7.

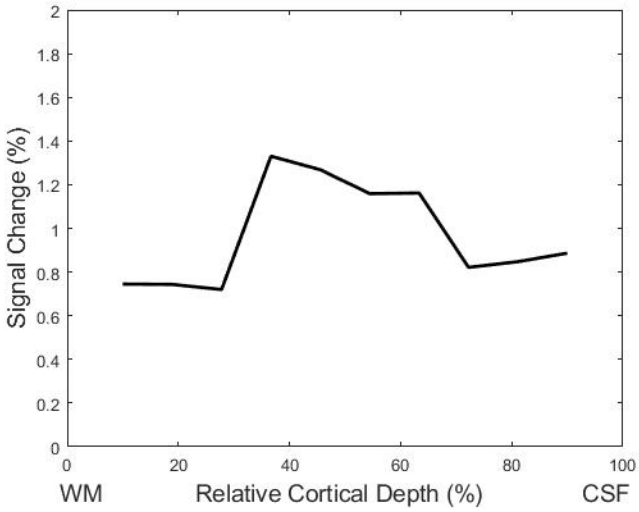


Figure S7: The simulated VASO response with 250μm resolution and no smoothing. Note the two potential peaks correspond to higher local CBV change in both side of stria of Gennari that can possibly be observed with higher acquisition resolution.

#### 1.6 Baseline CBV of the Vascular Compartments in the Cortical Vascular Model

Table S1 summarizes the baseline CBV of the vascular compartments in the cortical vascular model assuming a depth-dependent baseline CBV across the cortical depth. The first row shows the percentage of the baseline CBV in each vascular compartment, and the second row indicate the percentage of the baseline CBV relative to the total baseline CBV in the modelled region. The arterial and venous CBV fraction in the model are 43% and 57% respectively, that are in agreement with the reported values in the literature (Ito et al., 2005; Ito et al., 2001).

Table S1: The average values of the baseline CBV of the vascular compartments in the cortical vascular model. First row shows the percentage of the baseline CBV of each compartment, and the second row shows these values relative to the total baseline CBV.

| ***Vasculature Compartments*** | ***Average Baseline CBV (%)*** | ***Average Baseline CBV (%) [Relative to total CBV]*** |
|---|---|---|
| ***Laminar network*** | 2.2 % | 59 % |
| ***ICAs*** | 0.6 % | 16 % |
| ***ICVs*** | 0.8 % | 21 % |
| ***ICAs + arterioles*** | 1.6 % | 43 % |
| ***ICVs+ venules + capillaries*** | 2 % | 57 % |

#### 1.7 The diameters of the intra-cortical vessels corresponding to the constant baseline CBV scenario

The average diameter of the intra-cortical vessels (ICAs and ICVs) corresponding to the constant baseline CBV scenario are summarised in Table S1. These values have been calculated at each cortical depth using the cortical vascular model (Markuerkiaga et al., 2016), and are comparable with the values reported in a post-mortem human brain study by Duvernoy et al. (1981).

Table S2: The average diameter (in µm) of intra-cortical arteries and veins in the modelled vascular unit correspond to the constant baseline CBV scenario (section 2.4)

| ***The Average Diameter (µm) of the Intracortical Vessels*** | | | | | | | | |
|---|---|---|---|---|---|---|---|---|
| ***Vessel Type*** | **V4** | **V3** | **V2** | **V1** | **A4** | **A3** | **A2** | **A1** |
| ***Layer I*** | 69 | 53 | 34 | 21 | 43 | 34 | 21 | 13 |
| ***Layer II/III*** | 67 | 51 | 27 | | 42 | 32 | 17 | |
| ***Layer IV*** | 62 | 40 | | | 39 | 25 | | |
| ***Layer V*** | 55 | | | | 35 | | | |
| ***Layer VI*** | 43 | | | | 27 | | | |

#### 1.8 VASO and BOLD Signal Compartments

Figure S8 shows the simulated nulled (uncorrected) and BOLD-corrected VASO signals along with the extra- and intra-vascular BOLD signals corresponding to the best fit in the scenario shown in Figure 2. The BOLD contamination in the uncorrected VASO manifests as a signal increase near the cortical surface. When correcting for the $T_2^*$ dependency, the signal change peaks in the middle layers. The small intra-vascular BOLD signal change stems from the more oxygenated blood vessels, i.e., ICAs, arterioles and capillaries, and the extra-vascular BOLD signal increase near the surface is due to the dephasing of the tissue signal around venules and ICVs.

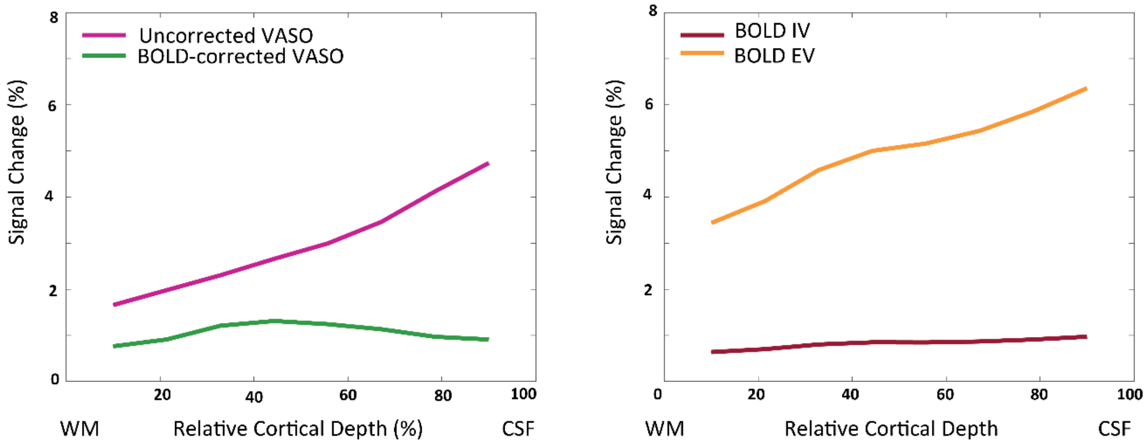


Figure S8: The uncorrected and BOLD-corrected simulated VASO profiles corresponding to the best fit (left). The Intra- and extra-vascular simulated BOLD signal correspond to the best fit (right). Both correspond to the scenario shown in Figure 2 of the manuscript.

### 2 References


Duvernoy, H.M., Delon, S., Vannson, J., 1981. Cortical blood vessels of the human brain. Brain Research Bulletin. 7, 519-579. https://doi.org/10.1016/0361-9230(81)90007-1.

Huber, L., Goense, J., Kennerley, A.J., Trampel, R., Guidi, M., Reimer, E., Ivanov, D., Neef, N., Gauthier, C.J., Turner, R., 2015. Cortical lamina-dependent blood volume changes in human brain at 7 T. Neuroimage. 107, 23-33. https://doi.org/10.1016/j.neuroimage.2014.11.046.

Ito, H., Ibaraki, M., Kanno, I., Fukuda, H., Miura, S., 2005. Changes in the arterial fraction of human cerebral blood volume during hypercapnia and hypocapnia measured by positron emission tomography. Journal of Cerebral Blood Flow & Metabolism. 25, 852-857. https://doi.org/10.1038%2Fsj.jcbfm.9600076.

Ito, H., Kanno, I., Iida, H., Hatazawa, J., Shimosegawa, E., Tamura, H., Okudera, T., 2001. Arterial fraction of cerebral blood volume in humans measured by positron emission tomography. Annals of Nuclear Medicine. 15, 111-116. https://doi.org/10.1007/BF02988600.

Markuerkiaga, I., Barth, M., Norris, D.G., 2016. A cortical vascular model for examining the specificity of the laminar BOLD signal. Neuroimage. 132, 491-498. https://doi.org/10.1016/j.neuroimage.2016.02.073.
